# Decoding shared versus divergent transcriptomic signatures across cortico-amygdala circuitry in PTSD and depressive disorders

**DOI:** 10.1101/2021.01.12.426438

**Authors:** Andrew E. Jaffe, Ran Tao, Matthew N. Tran, Stephanie C. Page, Kristen R. Maynard, Elizabeth A. Pattie, Claudia V. Nguyen, Amy Deep-Soboslay, Rahul Bharadwaj, Keith A. Young, Matthew J. Friedman, Douglas E. Williamson, Traumatic Stress Brain Research Group, Joo Heon Shin, Thomas M. Hyde, Keri Martinowich, Joel E. Kleinman

**Author notes:** Co-corresponding authors, and; 855 N Wolfe St., Ste 300; Baltimore, MD 21205.

## Abstract

Post-traumatic stress disorder (PTSD) is a debilitating neuropsychiatric disease with a projected lifetime risk of 8.7%. PTSD is highly comorbid with depressive disorders including major depressive disorder (MDD) and bipolar disorder (BD). It is hypothesized that the overlap in symptoms stems from partially shared underlying neurobiological mechanisms. To better understand shared and unique transcriptional patterns of PTSD and MDD we performed RNA-sequencing in the postmortem brain of two prefrontal cortex (PFC) regions and two amygdala (AMY) regions, from neurotypical donors (N=109) as well as donors with diagnoses of PTSD (N=107) or MDD (N=109) across 1285 RNA-seq samples. We identified a small number of differentially expressed genes (DEGs) specific to PTSD, mostly in the cortex compared to amygdala. PTSD-specific DEGs were preferentially enriched in cortistatin-expressing cells, a subpopulation of somatostatin interneurons. These PTSD DEGs also showed strong enrichment for gene sets associated with immune-related pathways and microglia, largely driven by decreased expression of these genes in PTSD donors. While we identified a greater number of DEGs for MDD, there were only a few that were specific to MDD as they showed high overlap with PTSD DEGs. Finally, we used weighted gene co-expression network analysis (WGCNA) as an orthogonal approach to confirm the observed cellular and molecular associations. These findings highlight the sub-population of cortistatin-expressing interneurons as having potential functional significance in PTSD and provide supporting evidence for dysregulated neuroinflammation and immune signaling in MDD and PTSD pathophysiology.

## Introduction

Post-traumatic stress disorder (PTSD) is a prevalent, debilitating disorder that develops in a subset of individuals following exposure to trauma. PTSD is highly comorbid with depressive as well as other mental health disorders (1–3). Indeed, epidemiological estimates suggest that >50% of individuals with PTSD also have a major depressive disorder (MDD) diagnosis (4–6). In addition to increased comorbidity with MDD, PTSD prevalence among individuals with a diagnosis of bipolar disorder (BD) is estimated to be 2-3 times that of the general population (7; 8). While PTSD is characterized by a unique set of clinical phenotypes, the disorder shares some diagnostic symptoms with MDD and BD, which also share overlapping symptom profiles. Some evidence suggests that comorbidity stems from shared mechanistic underpinnings, including overlapping genetic heritability as well as common environmental risk factors such as exposure to chronic stress and trauma(4). However, the cellular and molecular mechanisms that are unique to PTSD versus overlapping with MDD and BD are less well-understood.

Advances in imaging and recording technologies in humans and animal models have contributed to a greater appreciation of neuropsychiatric disorders, including PTSD, as disorders of brain circuits (9; 10). Functional magnetic resonance imaging studies in individuals with PTSD coupled with in vivo imaging and electrophysiological studies in animal models of behavior that are relevant for PTSD identified aberrant activity in neural circuits that encompass the amygdala and frontal cortical regions (10–12). These brain regions are critical for emotional regulation, including the expression and extinction of fear, functions which are frequently dysregulated in PTSD. Disruption of neural activity in these cortico-amygdala circuits is thought to reflect altered functional connectivity between the two regions, influencing behaviors that are frequently disrupted in PTSD including fear processing, anxiety and stress coping (^13–16^). In accordance with human neuroimaging studies that show altered signaling in cortico-amygdala circuits in PTSD (13; 16–18), studies in animal models of behavior demonstrate that function in these circuits is strongly impacted by exposure to stress and trauma. Moreover, experimentally manipulating neuronal activity in cortico-amygdala circuits or altering key cell signaling pathways in these brain regions is associated with changes in fear processing and anxiety (19–22).

At the cellular level, deficits in inhibitory neurotransmission in both frontal cortical regions and the amygdala have been implicated in depressive disorders and PTSD (23; 24). It is hypothesized that exposure to stress and trauma leads to deficits in inhibitory interneuron function, which impacts the balance of excitation and inhibition to alter neural activity and connectivity in cortico-amygdala circuits (25; 26). Moreover, there is strong evidence that GABAergic interneurons control neural activity and synaptic plasticity in these cortico-amygdala circuits to regulate fear-related behaviors in preclinical fear-conditioning models that are relevant for PTSD (27). However, how the molecular sequelae that follows exposure to stress and trauma impacts the function of specific interneuron cell types is not well understood, and how impaired inhibitory neuron function affects microcircuits that control connectivity between the amygdala and frontal cortical regions is not yet fully elucidated (25; 26; 28).

Immune- and inflammation-related signaling have more recently emerged as contributors to the development of PTSD as well as depressive disorders (29–32). While inflammatory markers and genes related to immune signaling have been reported to be altered in PTSD (33–35), findings indicate potentially complex changes, and many questions remain about the roles of these systems in disease. For example, whether observed changes result from central versus peripheral immune signaling pathways, and whether they reflect increased risk for PTSD, or epiphenomenon related to the pathophysiological sequelae of PTSD is not clear.

## Methods

### Human brain tissue

Postmortem human brains were donated through US medical examiners’ offices at the time of autopsy (total N=326), including the Office of the Chief Medical Examiner of: the State of Maryland (n=279), the District of Columbia (n=6), and of Virginia, Northern District (n=16), Western Michigan University Homer Stryker MD School of Medicine, Department of Pathology (n=24), and University of North Dakota Forensic Pathology Practice Center, Grand Forks County Coroner’s Office (n=1). Legal next-of-kin gave informed consent to brain donation according to protocols Maryland Department of Health and Mental Hygiene (MDHMH) # 12-24 (MD), National Institute of Mental Health (NIMH)# 90-M-014 (District of ColumbiaC and VirginiaA), and Western Institutional Review Board (WIRB) # 1126332 (MarylandD, Western Michigan UniversityMU, University of North DakotaND), respectively. Every brain received both a macroscopic and microscopic neuropathological examination at the time of autopsy by a board-certified neuropathologist. Brains were excluded from this study if there was evidence of cerebrovascular accidents, neuritic pathology, or other significant trauma to the brain that precluded it from further study.

The research conducted in this study was not considered human subjects research as defined by the HHS, as according to 45 CFR 46.102(f), a human subject is defined as a living individual about whom an investigator conducting research obtains data through intervention or interaction with the individual or identifiable private information.This research involved the analysis of RNA from postmortem human tissue.

A retrospective clinical diagnostic review was conducted on every brain donor, consisting of the telephone screening, macroscopic and microscopic neuropathological examinations, autopsy and forensic investigative data, two sources of toxicology data, extensive psychiatric treatment, substance abuse treatment, and medical record reviews, and whenever possible, family informant interviews (i.e., next-of-kin could be recontacted and was agreeable to phone contact,, which included the PTSD Checklist (i.e., PCL-5 and/or the MINI). A history of traumatic exposure including exposure to military combat, physical abuse, sexual abuse, emotional abuse, and/or other traumas were obtained as part of the telephone screening, records reviews, and/or PCL-5. A board-certified psychiatrist with expertise in PTSD reviewed every case in this study to rate presence/absence of PTSD symptoms. A summary of the trauma information is in Table S3.

All data were compiled into a comprehensive psychiatric narrative summary that was reviewed by two board-certified psychiatrists in order to arrive at lifetime DSM-5 psychiatric diagnoses (including substance use disorders/intoxication) and medical diagnoses. Non-psychiatric healthy controls were free from psychiatric and substance use diagnoses, and their toxicological data was negative for drugs of abuse. Every brain donor had either a medical examiner toxicological analysis, which typically covered ethanol and volatiles, opiates, cocaine and metabolites, amphetamines, and benzodiazepines. Every donor also received supplemental directed toxicological analysis using National Medical Services, Inc., including nicotine/cotinine testing, cannabis testing, and the expanded forensic panel in postmortem blood (or, in rare cases, in postmortem cerebellar tissue) in order to cover any substances not tested. The presence of opioids was determined by toxicology evaluations, including: codeine, morphine, oxycodone, hydrocodone, oxymorphone, hydromorphone, methadone, propoxyphene, fentanyl, 6-acetylmorphine (an active metabolite of heroin), and tramadol. If the medical examiner specifically noted the presence of any other opioids, then the subject was also included in this count.

### Tissue dissections

#### dACC

The dorsal anterior cingulate gyrus was identified visually on 1 cm thick coronal slab, on the mesial surface of the frontal lobe at the level of the genu of the corpus callosum. Gray matter from the cortical ribbon of the dACC, which was identified as the gyrus immediately dorsal to the corpus callosum, was dissected from the slab using a hand-held dental drill while the slab was positioned on dry ice after being removed from storage in a –80 C freezer.

#### dlPFC

Under direct visual guidance using a hand held dental drill, for the dorsolateral prefrontal cortex (dlPFC) dissections, grey matter tissue from the cortical ribbon was dissected from the crown of the middle frontal gyrus, from the coronal slab immediately anterior to the genu of the corpus callosum. Subcortical white matter was carefully trimmed from the area immediately below the middle frontal gyrus.

#### Amygdala

The medial and baso-lateral amygdaloid nuclei were dissected from the mesial superior temporal lobe for 0.75-1 cm thick coronal slabs of frozen human brains on dry ice using a stainless steel punch (8mm diameter Thermo Fisher Scientific Integra Miltex standard biopsy punch). Amygdaloid nuclei were dissected at the level of the anterior commissure, anterior thalamus and lentiform nucleus, which corresponded to the middle part of the amygdala along its anterior to posterior axis (36; 37). Tissue punch weights were in the range of 100-150 mg and were further powdered in frozen state before downstream extractions to ensure minimal sampling variability across all subjects in the study.

### RNA sequencing

Total RNA was extracted from all 1304 tissue samples using AllPrep DNA/RNA Mini Kit (Qiagen Cat No./ID: 80204) to concurrently extract RNA and DNA from the same piece of homogenized tissue in brain region- and diagnostic group-balanced batches of 96 samples. Paired-end strand-specific sequencing libraries were prepared from 300ng total RNA using the TruSeq Stranded Total RNA Library Preparation kit with Ribo-Zero Gold ribosomal RNA depletion which removes rRNA and mtRNA. An equivalent amount of synthetic External RNA Controls Consortium (ERCC) RNA Mix 1 (Thermo Fisher Scientific) was spiked into each sample for quality control purposes. RNA-seq cDNA libraries were genotyped with qPCR across 33 SNPs to establish sample identities with a genotype barcode. The libraries were then sequenced on an Illumina HiSeq 3000 at the LIBD Sequencing Facility, producing a median of 131.3 million (IQR: 115.4-146.3) fragments (across 100-bp paired-end reads) per sample.

### RNA-seq processing pipeline

Raw sequencing reads were processed using the same pipeline described in detail in Collado-Torres et al (38). Briefly, paired-end reads were mapped to the hg38/ GRCh38 human reference genome with splice-aware aligner HISAT2 version 2.0.4 (39). Feature-level quantification based on GENCODE release 25 (GRCh38.p7) annotation was run on aligned reads using featureCounts (subread version 1.5.0-p3) (40) with a median 43.8% (IQR: 37.3%-49.0%) of mapped reads assigned to genes. Exon-exon junction counts were extracted from the BAM files using regtools v.0.1.0 (41) and the bed_to_juncs program from TopHat2 (42) to retain the number of supporting reads (in addition to returning the coordinates of the spliced sequence, rather than the maximum fragment range) as described in Jaffe et al. (43). Annotated transcripts were quantified with Salmon version 0.7.2 (44) and the synthetic ERCC transcripts were quantified with Kallisto version 0.43.0 (45). For an additional QC check of sample labeling, variant calling on 740 common missense SNVs (containing the above 33 cDNA-genotyped SNPs) was performed on each sample using bcftools version 1.2. We generated strand-specific base-pair coverage BigWig files for each sample using bam2wig.py version 2.6.4 from RSeQC (46) and wigToBigWig version 4 from UCSC tools (47) for quality surrogate variable analysis (48) (as described below).

### Genotype data processing

Genotype data were processed and imputed as previously described (43). Briefly, genotype imputation was performed on high-quality observed genotypes (removing low quality and rare variants) using the prephasing/imputation stepwise approach implemented in IMPUTE2 (49) and Shape-IT (50), with the imputation reference set from the full 1000 Human Genomes Project Phase 3 dataset (51), separately by Illumina platform using genome build hg19. There were a total of 5 imputed batches in the current study, with 4 of 5 batches using an Illumina Infinium Omni2.5-8 kit (versions 1.2 or 1.3), and the remaining batch of 37 samples using the Infinium Omni5-4 kit. Imputed genotypes were merged across imputation runs/batches in the Oxford file format as dosages, then converted to plink file format as “hard call” genotypes (treating variants with posterior probabilities < 0.9 as missing). We retained common variants (MAF > 5%) that were present in the majority of samples (missingness < 10%) and that were in Hardy Weinberg equilibrium (at p > 1×10^-6^) using the Plink tool kit version 1.90b3a (52). Multidimensional scaling (MDS) was performed on autosomal LD-independent SNPs (variation inflation factor = 1.25, corresponding to R^2^ < 0.2) to construct genomic ancestry components on each sample, which can be interpreted as quantitative levels of ethnicity – the first component separated the Caucasian and African American samples, for inclusion as potential confounders in the differential expression analyses described below. The same 740 observed and imputed DNA-genotyped SNPs (as described above in the RNA-seq data processing) were further extracted across the 326 unique donors.

### Quality control and sample filtering

After completing the preprocessing pipeline on 1304 RNA-seq samples across 326 donors, we performed quality control assessments, including for sample identities and RNA-seq data quality. We first computed the pairwise genotype correlations across the 1304 RNA-seq samples among 235/740 high quality and moderate coverage variants (mean depth between 5-80, biallelic variants, and variant distance bias - VDB - p-values greater than 0.1). From this 1304 RNA x 1304 RNA correlation matrix, we expected clusters of four samples per donor, with high correlations among the 4 samples from the same donor and low correlations to all other samples. We subsequently computed the correlation between the 231/235 high-quality variants also present in the DNA-derived genotype data across these 326 donors (forming a 1304 RNA x 326 DNA correlation matrix). Here we expected each RNA sample to match DNA from its labeled donor. These two correlation matrices allowed us to a) identify and b) potentially recover sample identities, and comparison to the 33 cDNA genotypes further refined the processing steps where sample swaps occurred. Overall, 19 RNA-seq samples were dropped due to sample mis-identity issues, including 5 samples with sample contamination (two genomes in the RNA-seq library) and 11 samples with genotyping issues inconsistent with the study design (i.e. a fifth sample from the same donor with one region repeated) and 3 samples first identified in the cDNA libraries and confirmed in the RNA-seq libraries. The 5 samples with contaminations were identified by moderate correlations to DNA genotypes (∼0.4-0.6) and were indicative of pipetting issues during library preparations resulting in library mixing, as examinations of these mixed samples were always in adjacent wells on the 96 well preparation plates. There were 8 additional samples (from 4 pairs) that were identified as pairwise sample swaps that were reversed, and one sample with a mislabeled brain number. Our final sample characterizations and analyses were therefore performed on 1285 RNA-seq samples.

We then examined the distribution of sequencing and RNA quality metrics across group-region pairs, flow cells, and processing plates (Figure S1, Figure S11). ERCC spike-ins were uniformly distributed across brain regions and diagnosis groups (all p > 0.01), while metrics related to RNA quality (exonic mapping rate, mitochondrial mapping rate, RNA integrity number, and genome alignment rate) varied by brain region (all ANOVA p < 1e-4) but not diagnosis (all ANOVA p > 0.01). Examination of analogous effects by processing plate showed differences in ERCC spike-ins between the first 6 compared to the next 8 plates (Figure S11A) and lower RNA quality for plates 3 and 13 (Figure S11B). As plates were balanced by the primary outcomes of interest (region and diagnosis), we retained these samples in all downstream analyses and subsequently adjusted for these variables in differential expression analyses to reduce variation attributable to technical factors.

### Feature filtering by expression levels

We filtered lowly expressed features across all 1285 samples prior to expression analyses within and across brain regions. We calculated reads per kilobase per million (RPKM) genes (or exons) assigned during counting for genes and exons, and retained count data from 26,020 genes and 415,709 exons with RPKM > 0.2. Here we explicitly used the total number of gene or exon counts, and explicitly not the total number of aligned reads (which is sometimes used in RPKM calculations). We normalized exon-exon splice junction reads by scaling counts to 10 million reads with splice junctions (RP10M, analogous to total assigned gene counts for RPKM calculations). We used 10 million instead of 1 million both because it leads to easier visualization and because 10 million reads was the approximate number of spliced reads in an average library; this retained 217851 splice junction counts with RP10M > 0.75 that were at least partially annotated to one exon (ie annotated, exonic skipping or shifted exonic boundary junction classes). We lastly filtered pseudo-aligned normalized transcript counts (TPMs) using a cutoff of 0.2, leaving 101,515 transcripts for analysis.

### Degradation data processing for qSVA

We calculated quality surrogate variables to account for potentially latent RNA quality confounding (48). Here we used already-available RNA-seq degradation profiles from dlPFC [neuron 2019] (20 samples: 5 brains and 4 time points) and bulk amygdala [zandi preprint] (20 samples: 5 brains and 4 time points), and implemented a combined-region approach akin to Collado-Torres et al 2019. We therefore calculated mean coverage separately by strand across all 40 combined samples to define expressed regions with greater than 5 normalized reads and greater than 50bp (53). We then fit a linear model each expression region as a function of degradation time adjusting for brain region and donor as fixed effects. We then ranked the expressed regions by the degradation effect and created an input bed file with the top 1000 degradation-susceptible regions for coverage-based quantification in the 1285 post-QC RNA-seq samples described above. Subsequently quality surrogate variables (qSVS) for each sample were calculated once for the entire project from the top k principal components (PCs) of the expression in these 1000 degradation regions across all 1285 samples. We selected k = 19 using the BE algorithm (54) with the sva Bioconductor package (55).

### Differential expression analyses

We performed differential expression analyses across several different subsets of samples, for several different statistical models, at four feature summarizations. Our main analyses involved dividing this dataset into to regional groups: combining the two cortical regions (dlPFC + dACC) into a “cortex” dataset (N=641) and the two amygdala subregions (BLA + MedAmy) in an “amygdala” dataset (N=644). These analyses involved three groups of samples: neurotypical controls (“CONT”, patients with major depression (“MDD”) and patients with PTSD (“PTSD”), where the CONT group was generally the reference group in downstream regression analyses. Within each of these datasets, we fit one main statistical model that involved jointly estimating the effects of (a) PTSD vs CONT, (b) MDD vs CONT, (c) PTSD vs MDD, and (d) PTSD vs MDD vs CONT of the form:

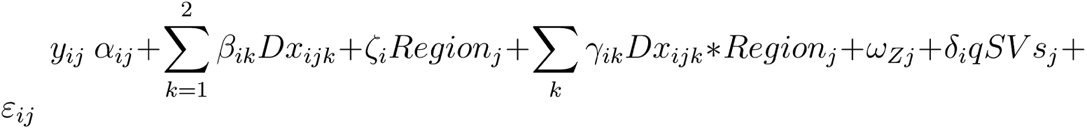

Where *y_ij_* are voom-normalized feature counts (on the log2 scale) (56) for feature *i* and sample, *β_i_*_1_is the log_2_ fold change for PTSD vs Control and *β_i_*_2_ is the log_2 fold change for MDD vs Control. We adjusted for subregions in the cortex (ie dACC vs dlPFC) and amygdala (ie BLA vs MedAmy) analyses, as well as the statistical interaction between the subregion term and each diagnosis main effect term. We further adjusted for the vector of fixed effects potential observed confounders “Z”, including age, sex, chrM mapping rate, rRNA rate, exonic mapping rate, RIN, overall mapping rate, the ERCC bias factor (e.g. root mean square error) and then quantitative ancestry factors 1,2,3, 8, 9, and 10 (which were all associated with diagnosis groups). We further adjusted for the vector of fixed effects latent confounders “qSVs” (specifically 19 qSVs, described above). Here, as there were multiple regions from the same donors, *α_ij_* was parameterized as a random intercept using the ‘duplicateCorrelation’ function in limma, with donor as the blocking variable. We therefore used linear mixed effects modeling (rather than regular linear regression) to fit the above model, once to the “cortex” dataset, and again to the “amygdala” dataset. PTSD vs CONT, MDD vs CONT, and PTSD vs MDD effects were converted to empirical Bayes-moderated T-statistics, with corresponding p-values, and Benjamani-Hochberg-adjusted (BH-adjusted) p-values using the limma topTable function. We also used the topTable function to calculate an F-statistic to test mean differences between the three diagnosis groups from *β_ik_*, with corresponding p-values and BH-adjusted p-values. We fit secondary models to the “cortex” and “amygdala” datasets recoding the diagnosis group variable into a binary “PTSD-only” variable, comparing patients with PTSD (coded as 1) to a combined MDD and Control group (coded as 0) to estimate PTSD-specific effects with the exact same adjustment terms as the standard three-level diagnosis variable. We further fit secondary models to the “Joint”/combined dataset of all 1285 samples with linear mixed effects modeling, where there were 3 region terms (instead of 1 above) and 6 region-by-diagnosis interaction terms (instead of 2 above) which were used to calculate overall diagnosis effects across all regions as well as an F-statistic per model for overall diagnosis-by-region interaction effects. We lastly fit secondary models within each subregion using linear regression (since there were no repeated donors within each region) and dropped main effect and interaction terms related to region from the above model (ie \zeta and \gamma) to estimate the effects of diagnosis within each region. We fit these models separately at the feature levels of genes, exons, junctions, and transcripts. As we used TPMs rather than raw counts for transcript-level analyses, we skipped the voom step.

### Gene Ontology and gene set enrichment analyses

Unless otherwise noted, we used the compareCluster() function from clusterProfiler (57) version 3.14.3 for gene ontology (58; 59) and KEGG (60) enrichment analyses with the set of Ensembl gene IDs expressed in genes.

### WGCNA

We performed weighted gene co-expression network analysis (WGCNA) using the WGCNA R package (61) (package version 1.69 and R version 3.6.1). WGCNA analyses were performed in the four brain regions separately as well as in the two broad region groups (amygdala and cortex) and the full dataset, resulting in 7 total WGCNA analyses. After filtering out lowly expressed genes (cutoff mean RPKM > 0.2), the log_2_(RPKM+1) normalized expression values were “cleaned” using the cleaningY function from the jaffelab R package (62) (version 0.99.30). Specifically, the same covariates as modeled above were regressed out of the expression matrix: mitochondrial RNA rate; rRNA rate; gene assignment rate; RIN; mapping rate; ERCC spike-in error; genomic ancestry components 1, 2, 3, 8, 9, and 10; and latent quality surrogate variables 1-19, while preserving the effects of diagnosis, age and sex, and, when applicable, brain region and their interaction (in the combined-region analyses). WGCNA was run using the same strategy for each of the seven runs: automated determination of the soft thresholding parameter (using the ‘pickSoftThreshold’ function), and then constructing co-expressed modules (using the ‘blockwiseModules’ function) using signed networks with “bicor” correlation, with a minimum module size of 30, mergeCutHeights of 0.25, and no reassignment. We applied gene ontology enrichment analysis (GO) using clusterProfiler (57) (version 3.14.3) to understand the biological enrichments of our clusters. We subsequently tested the association between each module eigengene and diagnosis, adjusting for age, sex for single-region analyses, and further adjusted for brain region, and the interaction between brain region and diagnosis, as well as random effects of donors, when performing multi-region analyses. Note we did not account for the other confounders as their effects were regressed out of the expression data prior to module construction.

### Sensitivity analyses

For each of the four brain regions, we performed sensitivity analyses for the following observed covariates sequentially: opioid use (based on toxicology), exposure to trauma (lifetime, based on narratives/medical history), antidepressant use (based on toxicology), antidepressant use (lifetime, based on narratives/medical history), and manner of death (natural, suicide, undetermined, accidental). For each sensitivity analysis - which considered a single confounder from the previous list, we adjusted the original model (above) to include each additional covariate, and then compared the original regression coefficients for each diagnosis (β) to the further-adjusted diagnosis coefficient (β^*c*^, such that we ran 5 different sensitivity analyses)

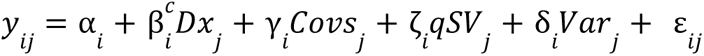

### Cell type enrichment analyses

We used functionality in the CSEA package to estimate cell type enrichments using pre-defined gene sets obtained from Mouse BAC-trap lines (63). We performed an analogous form of CSEA for human cell types using Fisher’s exact tests on pre-defined cell type-specific genes described in Tran et al (64).

### RNAscope single molecule fluorescence *in situ* hybridization

Postmortem dlPFC and BLA were dissected as previously described from two adult males with no known psychiatric illnesses. Brain tissue was equilibrated to −20°C in a cryostat (Leica, Wetzlar, Germany) and serial sections of dlPFC and BLA were collected at 10 μm. Sections were stored at −80°C until completion of the RNAscope assay.

We performed *in situ* hybridization with RNAscope technology utilizing the RNAscope Fluorescent Multiplex Kit V2 (Cat # 323120 Advanced Cell Diagnostics [ACD], Hayward, California) according to manufacturer’s instructions and as previously described (65). Briefly, tissue sections were fixed with a 10% neutral buffered formalin solution for 30 min at room temperature, series dehydrated in ethanol, and pretreated with hydrogen peroxide at RT for 10 minutes then with protease IV for 30 min. Sections were incubated with a custom-designed Channel 4 *CORT* probe (Cat # 593341-C4, Advanced Cell Diagnostics, Hayward, California) and commercially available *CRHBP*, *GAD2*, and *SST* probes (Cat #s 573411, 415691-C3, 310591-C2) for 2 hours and stored overnight in 4x SSC (saline-sodium citrate) buffer. Probes were fluorescently labeled with Opal dyes (PerkinElmer, Waltham, MA). Opal dyes were diluted at 1:500 and assigned to each probe as follows: Opal520 to *CRHBP*, Opal570 to *SST*, Opal620 to *CORT,* and Opal690 to *GAD2*). Confocal lambda stacks were acquired using a Zeiss LSM 780 equipped with a 63X/1.4NA objective, a GaAsP spectral detector and 405, 488, 561, and 647 lasers. All lambda stacks were acquired using the same laser intensities, linear unmixing performed as previously described, and images were processed with our dotdotdot software (65). Summarized nuclei/ROI-level and transcript/dot-level data were analyzed using Pearson correlation analyses in R.

## Results

### Cohort and sequencing data descriptions

We generated deep bulk/homogenate RNA-sequencing (RNA-seq) data from postmortem human tissue in two subregions of the frontal cortex (dorsolateral prefrontal cortex, dlPFC, and dorsal anterior cingulate cortex, dACC) and two subregions of the amygdala (basolateral amygdala, BLA, and medial amygdala, MeA; **Table S1**) from neurotypical donors versus donors with diagnoses of MDD, BD and/or PTSD to better understand shared and divergent gene expression. The MDD group consisted of 109 patients with a primary diagnosis of MDD (DSM, 5th edition(66)). The PTSD group consisted of a total of 107 patients with DSM-5 PTSD diagnosis: 78 patients had a primary PTSD diagnosis and 29 patients had a secondary PTSD diagnosis. Of those with a secondary PTSD diagnosis, 28 had a primary diagnosis of BD and 1 had a primary diagnosis of bipolar not otherwise specified (BpNOS). Types of trauma were characterized for all donors in the PTSD group - a majority were not exposed to combat (76.6%, **Table S2**), but rather had high rates of childhood maltreatment (66%, **Table S3**). Donors in the PTSD group also had high rates of comorbidity for MDD (62.6% with a secondary diagnosis, **Table S2**) and substance use disorder (SUD) (77.6% with a secondary diagnosis, **Table S2**). The neurotypical control group was composed of 109 donors.

After extensive and rigorous quality control of RNA-seq data (see Methods, **Figure S1A-B**), we performed differential expression and network analyses across 1285 samples from these 325 unique donors (**Table 1**). Gene expression analyses were performed on the same set of 26,020 expressed genes across all samples. We performed principal component analysis (PCA) of these data to better characterize global patterns of gene expression (**Figure S1C**). Top components of gene expression variation related to brain region and various measures of RNA quality (**Figures S1D**), with the first principal component (PC1, 20.4% of variance explained) associating strongly with RNA quality **(Figure S1E**), and distinguishing broad cortical and amygdala regions (with smaller differences within subregions, particularly within the amygdala (**Figure S1F**)). These data recapitulate established cytoarchitecture, as we observed greater local expression homogeneity among amygdala nuclei than cortical subregions, with greater amygdala similarity to the dACC than the dlPFC (67). We employed quality surrogate variable analysis (qSVA) (48), which defines degradation-susceptible genes across dlPFC and broad amygdala (see Methods), for downstream differential expression and network analyses to control for both observed and latent potential confounders.

**Table 1:** demographic and RNA quality information for the subjects and associated brain tissue in this study.

| **Group** | **Control** | **MDD** | **PTSD** |
| --- | --- | --- | --- |
| **Total subjects** | 109 | 109 | 107 |
| basolateral amygdala | 108 | 109 | 105 |
| medial amygdala | 109 | 106 | 107 |
| dorsal anterior cingulate cortex | 107 | 109 | 106 |
| dorsolateral prefrontal cortex | 107 | 108 | 104 |
| **Sex (% M)** | 78.9 | 55 | 50.5 |
| **Race (% Cauc)** | 69.7 | 80.7 | 87.9 |
| **Age: Mean (SD)** | 49.6 (15.1) | 45.9 (14.5) | 40.8 (11.3) |
| **RIN: Mean, all regions (SD)** | 7.4 (0.8) | 7.3 (0.9) | 7.3 (0.9) |
| BLA: Mean (SD) | 7.1 (0.8) | 6.9 (0.8) | 6.8 (0.7) |
| MeA: Mean (SD) | 7.0 (0.8) | 6.8 (0.7) | 7.2 (1.1) |
| dACC: Mean (SD) | 7.5 (0.8) | 7.5 (0.7) | 7.6 (0.7) |
| dIPFC: Mean (SD) | 7.9 (0.6) | 8.0 (0.9) | 7.7 (0.6) |
| **PMI (hr): Mean (SD)** | 29.4 (11.1) | 26.7 (7.71) | 29.1 (10.7) |
| **Smoke (%)** | 16.5 | 65.1 | 73.6 |
| **Opioids (%)** | 6.42 | 62.4 | 66.4 |
| **Manner of Death (% Suicide)** | NA | 22.9 | 24.3 |
| **Drug Related Death (%)** | NA | 45.9 | 69.2 |

### Expression differences related to PTSD diagnosis

We first explored the gene expression effects of PTSD diagnosis versus neurotypical control donors. Since subregions from the same broader regions showed more similar global expression patterns (**Figure S1F)**, we hypothesized that conducting primary analyses where subregions within the cortex and then the amygdala were combined to assess broader cortex and amygdala could increase statistical power to detect differentially expressed genes. We therefore identified the effects of PTSD versus neurotypical controls using linear mixed effects modeling within 641 broader cortex samples and then within 644 broader amygdala samples (including MDD samples, see Methods), allowing for differential PTSD versus neurotypical control effects within subregions (while simultaneously estimating MDD versus control effects, see Methods). We identified 41 such PTSD differentially expressed genes (DEGs) in cortex (**Figure 1A**) and 1 PTSD DEG in amygdala (**Figure 1B**) at genome-wide significance (FDR < 0.05), while a more liberal threshold of FDR < 0.1 identified an additional 78 genes in cortex (with no additional genes in amygdala). We highlight several representative differentially expressed genes in PTSD versus neurotypical control donors in cortex including decreased expression of *CORT*, a key marker of cortistatin-positive interneurons (68) (**Figure 1C**), and increased expression of the histone deacetylase *HDAC4* (**Figure 1D**), as well as *SPRED1,* which encodes a protein involved in the Ras/MAPK signaling pathway (**Figure 1E**). In amygdala, there was a single gene consistently downregulated in PTSD versus neurotypical controls across both subregions - *CRHBP* (**Figure 1F**), the gene encoding corticotropin-releasing hormone binding protein, which is an antagonist of the stress hormone corticotropin-releasing hormone (CRH) (69). Overall, cortical regions showed more association with PTSD than amygdala subregions, and the observed expression differences were largely consistent across subregions of the cortex, with only 5 genes showing marginal interaction (at p<0.01) between PTSD diagnosis and cortical subregion (*NRSN1*, *PHF20L1*, *RP11-505E24.2*, *OXLD1*, *CARD8-AS1*), and *CRHBP* only showing modest interaction between PTSD and amygdala subregions (p=0.037).

**Figure 1:**
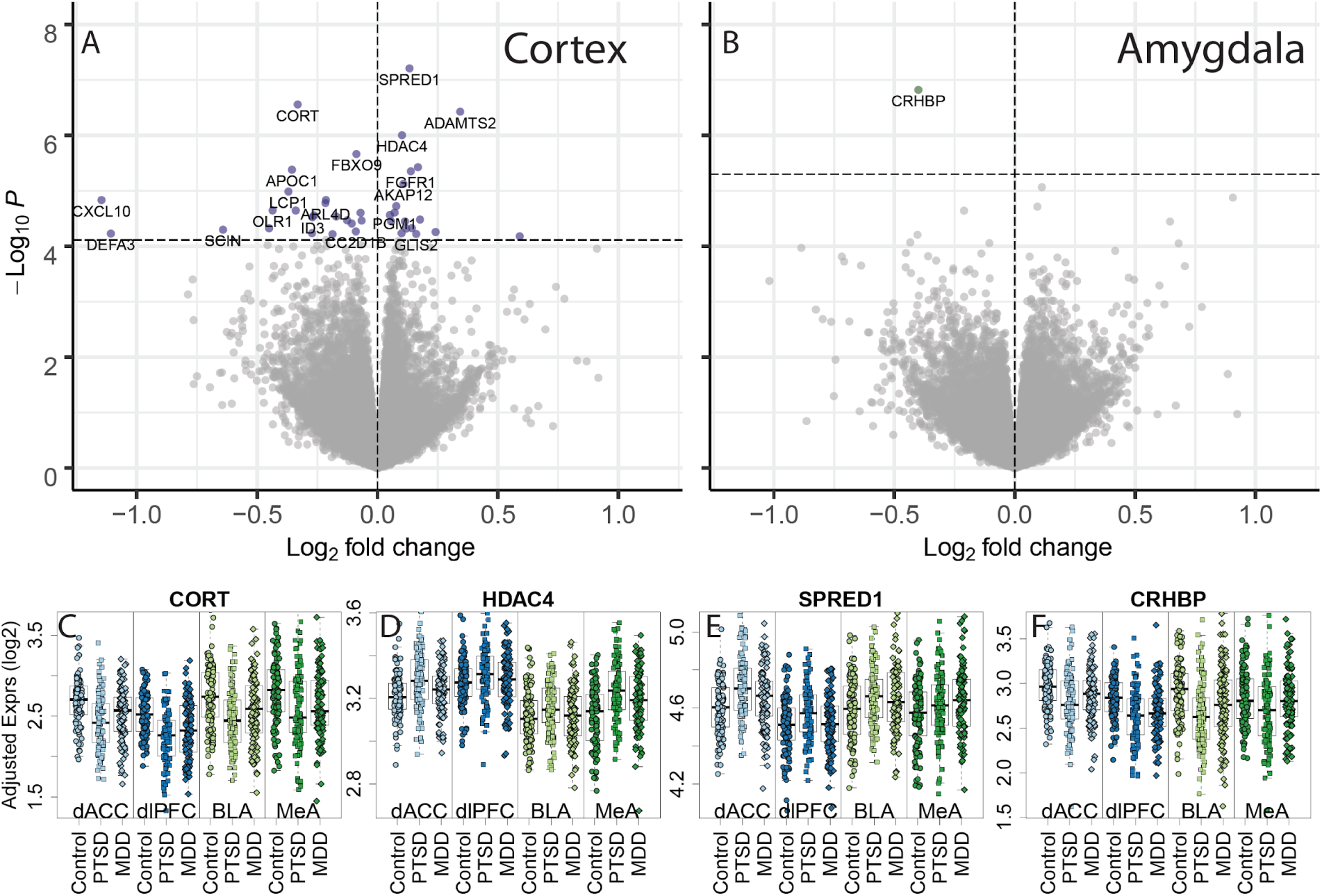
Differential gene expression associated with PTSD diagnosis, compared to neurotypical controls. Volcano plots for (A) cortex and (B) amygdala subregion-combined dataset. P-values were calculated using linear mixed effects modeling and the horizontal dashed line indicates the p-value that controls a false discovery rate (FDR) < 0.05. Positive log2 fold changes indicated higher expression in PTSD versus neurotypical subjects and negative log2 fold changes indicated lower expression in the PTSD group. Example differentially expressed genes include (C) CORT (D) HDAC4, (E) SPRED1, and (F) CRHBP, with “Adjusted” expression on the y-axis (regressing out unwanted technical and clinical confounders, preserving group and region effects, see Methods).

We next performed secondary analyses within each of the four subregions to identify additional DEGs associated with PTSD diagnosis. Differential expression statistics were highly correlated with the combined subregion analyses, with the cortical associations driven predominantly by dACC and the amygdala associations driven primarily by BLA (**Figure S2**). The cortical subregions again showed more PTSD DEGs, with 16 genes in the dACC (**Figure S3A**) and 1 gene in the dlPFC (**Figure S3B**) (and no genes in amygdala subregions) at genome-wide significance (FDR < 0.05). Using a more liberal cutoff of FDR < 0.1, we identified 74 unique genes across the cortical subregions (dACC: 72, dlPFC: 3, with one gene shared: *AC124804.1,* a novel transcript, antisense to *SDK2*) and 18 unique genes across amygdala subregions (BLA: 3, MeA: 16 with one gene: *CORT,* shared). Joint analysis of all data identified 117 genes with consistent PTSD versus control effects across all 4 subregions (at FDR < 0.05, with 276 genes at FDR < 0.1, **Figure S4**) further highlighting the similar effects of PTSD across multiple brain regions. Interestingly, these cross-region results were largely driven by the amygdala (predominantly BLA) and not cortex, even though the cortex had more DEGs when considered alone (**Figure S2**). A comprehensive list of all differential expression statistics for all expressed genes and all statistical models is presented in **Data S1**.

We further interrogated the role of potential confounders and risk factors across these PTSD associations to gene expression in a series of sensitivity analysis to determine the robustness of our differential expression model. We tested a series of additional potential variables (including treatment for antidepressants and presence of opioids via toxicology) for attenuating the DEGs identified above in each brain region. Overall, subsequently adjusting our models for these variables had minimal effects on our differential expression signal across all expressed genes, including those identified as DEGs (**Figure S5**). We further examined the role of sex on our identified DEGs using sex-specific analyses and found that subsets of our DEGs were more strongly explained by effects within a single sex (**Figure S6**). Lastly, we attempted to replicate our PTSD DEGs using results from a recent manuscript using a subset of the same donors (N=56) in the dACC (70). Across the 57 genes considered expressed in those data (of 72 identified here), 46 were directionally consistent (80.7%), and 9 were further genome-wide significant (15.8%, at FDR< 0.1), with highly correlated log2 fold changes across these 57 genes (=0.58, p=1.56e-6, **Figure S7A**). Analogous analyses using the 193 expressed DEGs in those data (among 196 identified) showed similar directional consistency (82.3%) and correlation (=0.56) of log_2_ fold changes in our data, but had a much lower overall replication rate (9/193, 4.7%). Another key difference between studies involved the inclusion of patients with BD in this study’s PTSD group. Sensitivity analyses in the dACC excluding the 28 donors in the PTSD group with a primary BD diagnosis yielded highly concordant DEGs as identified in the full dataset (via global t-statistic correlation, \rho = 0.963). Additional analyses comparing the PTSD (n=77) versus BD (n=28) primary diagnoses within the PTSD group showed little global correlation to the full PTSD versus control group effects (\rho = 0.11), and identified a single significant gene (*RPL13P12,* p=5.4e-8), which was not a PTSD (versus control) DEG (p=0.41). Taken together, these analyses identified robust sets of differentially expressed genes associated with PTSD, and which are unassociated with substance abuse and primary mood disorder diagnoses, in the largest postmortem brain dataset of PTSD to date.

### Gene sets and cell types associated with PTSD

We next performed gene set and pathway enrichment analyses to identify convergent biological and molecular functions associated with PTSD within and across brain subregions. Here we used more liberal significance thresholds to define PTSD DEGs (using marginal p < 0.005 rather than FDR control) and directionality to facilitate these analyses, and tested for enrichment among DEGs more highly and more lowly expressed in PTSD cases versus neurotypical controls. Overall, genes associated with PTSD showed the strongest enrichment for immune-related gene sets and pathways in both the cortex and amygdala (**Figure 2A**, **Table S4**), largely driven by decreased expression of genes in these pathways in patients with PTSD. Interrogating PTSD differences within subregions further identified unique molecular associations. For example, the MeA and dlPFC each showed decreased expression of genes associated with receptor ligand activity (that were further marginally significant in other regions). Interestingly, dlPFC associations were driven by eight genes (*CORT*/*CSF1*/*SST*/*OSTN*/*CXCL10*/*CXCL11*/*GDF9*/*CCL3*) and MeA associations by ten genes (*CORT*/*TNFSF10*/*CXCL11*/*SFRP2*/*OSGIN2*/*OGN*/*IGF2*/*CTF1*/*CCL5*/*TTR*) with only two genes in common (*CORT* and *CXCL11*), highlighting the convergence of molecular functions across brain regions.

**Figure 2:**
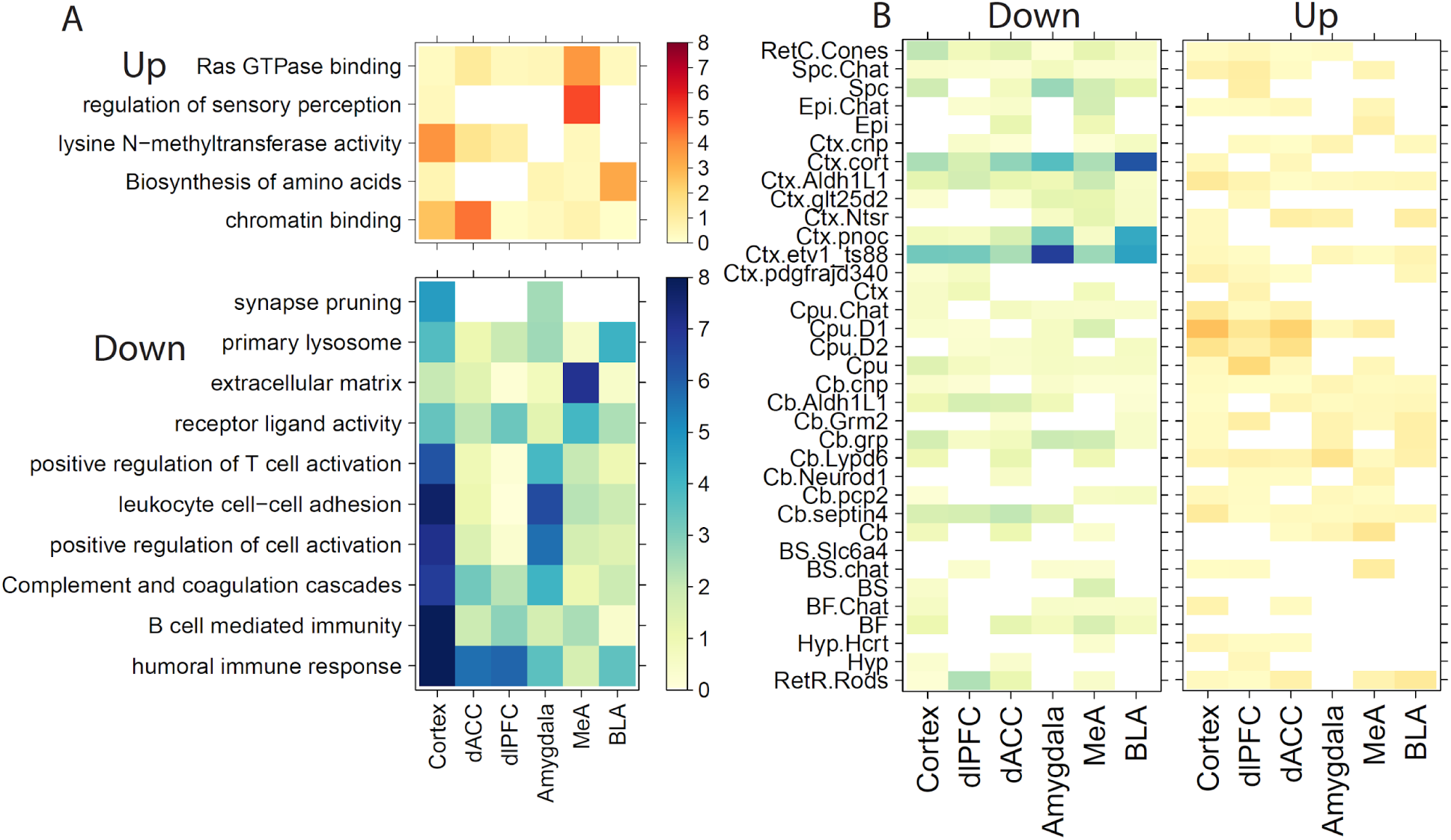
Molecular and cellular enrichments for genes associated with PTSD versus neurotypical controls. A) Gene set enrichment and (B) cell specific enrichment analyses for genes more highly expressed in PTSD (“Up”) or more lowly expressed (“Down”) compared to neurotypical donors. Color indicates-log10(p-values).

We next used cell type-specific enrichment analyses (CSEA) (63) to identify cell types that preferentially express these sets of differentially expressed genes. We used cell type-specific reference profiles generated with translating ribosomal affinity purification (TRAP) techniques from transgenic BACarray reporter mice (63; 71; 72) for these enrichment analyses because molecular annotations from the human brain are largely restricted to data obtained from single nucleus RNA sequencing (snRNA-seq), which lacks expression data from neuronal processes (73). We found consistent enrichment of cortistatin-positive interneurons (“Ctx.cort”) and immune cells (“Ctx.etv1_ts88”) among genes with increased expression in donors with PTSD compared to neurotypical controls. Stronger effects were observed in the amygdala, particularly the BLA, compared to the cortex (**Figure 2B**). We note that the reference cell type profile for Layer 5a corticostriatal interneurons (Ctx.etv1_ts88) was indicated as containing contamination with lymphoid cells (immune cells including microglia) (71; 72). These findings were robust to the choice of cellular specificity of input genes (specificity threshold/pSI ranges from 0.05 to 1e-4, **Table S5**) and convergent with the gene set enrichment analyses described above, particularly related to decreased expression of immune-related gene sets in PTSD. For example, using a specificity threshold of pSI < 0.01 and the BLA, immune cell enrichments were driven by decreased expression of *FERMT3*, *CRHBP*, *FOLR2*, *PTGS1*, *SLCO1C1*, *P2RY13* and *GLT8D2* (odds ratio, OR= 8.9, p=2.94e-5) and cortistatin-positive interneuron enrichments were driven by decreased expression of *NPY*, *CORT*, *CRHBP*, *DLL3*, *NXPH2*, and *SST* (OR=23.7, p=5.9e-7).

Since the above enrichment analyses used cell-type reference data from the mouse cortex it is unclear whether cell-type enrichments are conserved in the amygdala where cell composition may differ. We therefore sought to confirm immune-related expression enrichment using snRNA-seq data that we recently generated in the human amygdala (64). Cell types were defined using data-driven clustering, cell type-enriched genes were identified for each cluster, and enrichment testing was performed for genes differentially expressed in PTSD in amygdala and its subregions to each cluster. We again found preferential enrichment of immune-related genes - here represented by the microglia cluster (764 nuclei) - among those genes whose expression is decreased in donors with PTSD versus neurotypical controls. These enrichments were strongest in the combined amygdala analysis (OR=4.35, p=1.28e-16) driven by the BLA (OR=2.26, p=1.21e-05) without any significant enrichment in the MeA (p=0.42). As there were many thousands of genes associated with each cell type (see Methods), we performed secondary analyses taking the top 2000 genes associated with each cell type (to use the same number of genes for enrichment analyses), which strengthened our original observations (amygdala: OR=11.5,p=2.8e-28; BLA: OR=4.6, p=8.0e-9). We further only found marginal evidence for inhibitory interneuron enrichments associated with decreased expression in PTSD, particularly in the BLA (Inhib2: OR=2.87,p=3.7e-4; Inhib3: OR=2.67, p=1.1e-3). This negative result likely resulted from low levels of *CORT* gene expression in these snRNA-seq data (**Figure S8**), due to the low prevalence of cortistatin-expressing cells. Low expression levels of other genes driving enrichment of the cortistatin-expressing interneuron population also likely contributed to this effect (**Figure S8**), which is in contrast to the genes driving the immune enrichments, which are relatively much more highly expressed (**Figure S9**).

To validate the co-expression of our interneuron-related DEGs in amygdala and dlPFC we therefore used a complementary RNAscope single molecule fluorescence in situ hybridization (smFISH,see Methods) approach, which has increased sensitivity for rare cell populations compared to snRNA-seq. We targeted expression of *CORT*, *SST*, and *CRHBP* (DEGs for PTSD diagnosis in at least one brain region, **Data S1**), as well as *GAD2* (not a DEG) as a cell marker of inhibitory GABAergic cells. Given the low prevalence of *CORT* and *SST* transcripts, we captured images from locations in tissue sections of the BLA where *CORT* and/or *SST* were expressed (since the two fluorophores are difficult to distinguish by eye, **Figure 3A**) for a total of 489 nuclei/regions-of-interest (ROIs) across 32 images (see Methods). We compared expression levels of these genes across all nuclei, and generally found high correlations (**Figures 3B**), ranging from the highest correlations between *CORT* and *SST* (*ρ* = 0.72,p=8.7e-84) to the lowest between *GAD2* and *CRHBP* (*ρ* = 0.166, p=2.1e-13). Almost all *SST*+ interneurons co-expressed *CORT* (64/65 ROIs using 5 dot cutoff to classify as “expressed”) whereas only 64/183 *CORT*+ interneurons co-expressed *SST*. Our top amygdala DEG - *CRHBP* - showed co-expression with *CORT* and *GAD2* across many ROIs in both brain regions (74; 75). Together, these results support the hypothesis that specific subpopulations of GABAergic interneurons are functionally impacted in PTSD.

**Figure 3:**
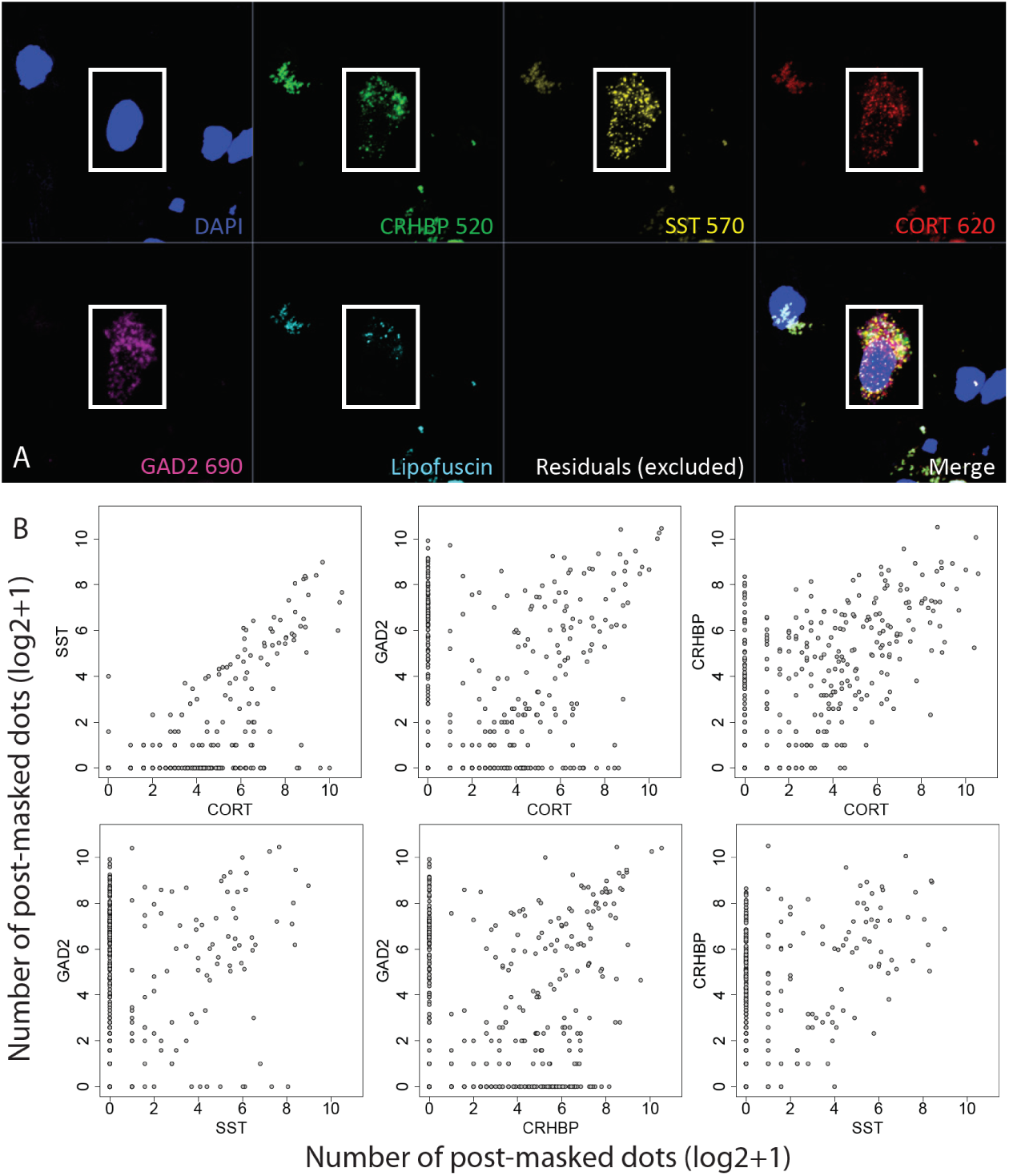
Single molecule fluorescence in situ hybridization (smFISH) validation of interneuron co-expression. (A) Representative image of co-expressing region-of-interest (ROI)/nucleus across multi-channel image. (B) Pairwise co-expression plots among four target genes, where axes indicate the number of post lipofuscin-masked segmented transcript dots (on the log2 scale)

**Figure 4:**
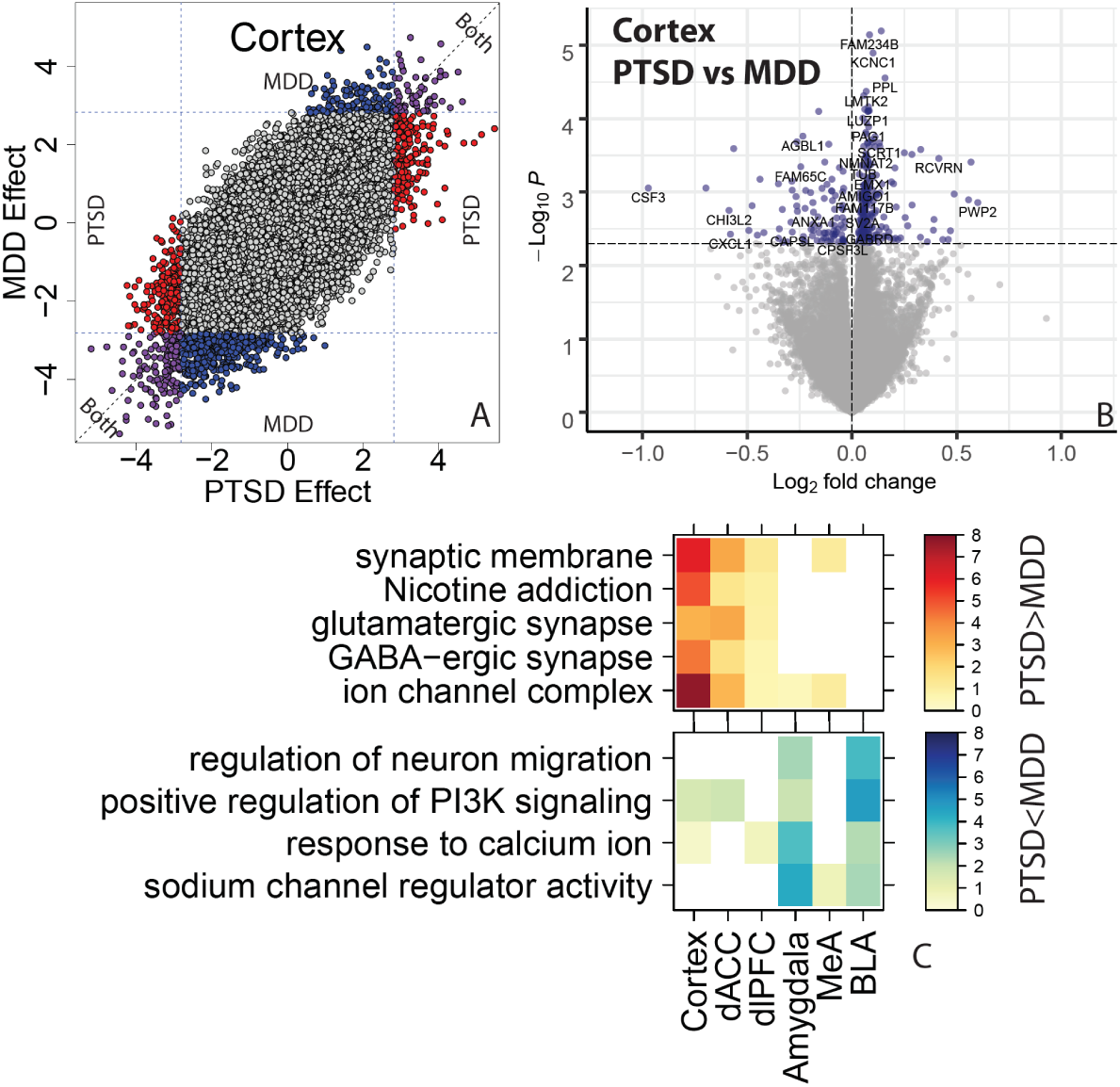
Contrasting PTSD and MDD effects on gene expression. (A) Scatterplot comparing the t-statistics for MDD versus control differential expression effects (y-axis) against PTSD versus control effects (x-axis). Colors indicate marginal significance at p < 0.005 for PTSD (red), MDD (blue), or both (purple). (B) Volcano plot directly comparing PTSD and MDD groups to each other. Horizontal line indicates marginal P < 0.005. (C) Gene set enrichment analyses for genes differentially expressed between PTSD and MDD, stratified by directionality.

### Untangling gene expression associations of PTSD and MDD diagnoses

We next incorporated RNA-seq data from MDD donors to better understand the gene expression differences unique to PTSD. We first compared patients with MDD to neurotypical controls among the broader cortical and amygdala brain regions, and again identified a larger number of differentially expressed genes in the cortex (182 genes at FDR < 0.05, 352 at FDR < 0.1, **Table 2**) compared to amygdala (0 genes at FDR < 0.05, 1 at FDR < 0.1). These cortical differences were driven by the dACC (249 genes at FDR <0.1) compared to dlPFC (2 genes at FDR<0.1) as in our PTSD effects. There were similarly increased MDD differences in the MeA (16 genes at FDR < 0.05, 32 at FDR < 0.1) and no differences in BLA when stratifying the amygdala into subregions. Genes with decreased expression in MDD donors compared to neurotypical controls showed analogous enrichment of immune-related processes in the cortex using both gene set enrichment analysis (**Table S6**) and CSEA (“Ctx.etv1_ts88” cell type, **Table S7**). Cell type enrichments related to cortistatin-positive interneurons were attenuated compared to PTSD effects, particularly in the amygdala (best p-value = 0.01)

**Table 2:** Number of differentially expressed genes in each dataset/brain region at two false discovery rate (FDR) cutoffs.

|  | **PTSD vs CONT** |  | **MDD vs CONT** |  |
| --- | --- | --- | --- | --- |
| **Dataset** | **FDR < 0.05** | **FDR < 0.1** | **FDR < 0.05** | **FDR < 0.1** |
| **Cortex** | 41 | 119 | 182 | 352 |
| **dACC** | 16 | 74 | 67 | 249 |
| **dIPFC** | 1 | 3 | 1 | 2 |
| **Amygdala** | 1 | 1 | 0 | 1 |
| **BLA** | 0 | 3 | 0 | 0 |
| **MeA** | 0 | 16 | 16 | 34 |
| **Joint** | 117 | 276 | 55 | 192 |

Globally, there was high concordance between PTSD and MDD effects on gene expression (**Figure S10**, *ρ* range=0.647-0.695), with highly overlapping differentially expressed genes at marginal significance in each brain region or subregion (all Fisher’s p-value < 1.72e-46). While global effects were correlated and significant genes overlapping, there were nevertheless variation among significantly differentially expressed genes across the two disorders. For example, among the genes marginally associated with MDD in each subregion, only a quarter were significantly differentially expressed comparing PTSD to controls (each at p < 0.005), and among those genes marginally associated with PTSD, only a third of genes in cortical regions and a quarter of genes in amygdala regions showed similar marginal association in MDD.

We therefore directly compared expression differences between PTSD and MDD donors to better partition these differences across diagnoses (see Methods). There were few genes that were genome-wide significantly different between these groups (at FDR< 0.1), with increased expression of *KCNC1*, *FAM234B* and *RASD2* and decreased expression of *CH507-513H4.4* in PTSD versus MDD in cortex, decreased expression of *LMCD1* in PTSD in MeA and decreased expression of *DNAH11* in PTSD in dlPFC. We then considered the sets of marginally significant (at p < 0.005) genes that were different between donors with PTSD compared to those with MDD. In cortex, genes more highly expressed in PTSD versus MDD were associated with neuronal processes and synapses (including both inhibitory and excitatory), whereas genes with decreased expression in PTSD versus MDD in amygdala were associated with neuronal migration and PI3K signaling (**Table S8**). There were no enrichments for the immune-related gene sets for these disorder-specific contrasts, suggesting decreased expression of immune processes and/or microglia involvement were shared across both disorders, relative to neurotypical individuals (**Table S9**). These results together suggest largely similar transcriptomic changes in PTSD and MDD compared to neurotypical donors.

### Co-expression analyses provide convergent evidence for immune and interneuron associations

We lastly performed two forms of co-expression analysis to better understand network-level gene expression differences between PTSD and MDD. First, we performed weighted gene co-expression analyses (WGCNA) at the broad region and then subregion levels, analogous to differential expression models above. This WGCNA approach assigns each gene to an individual module (“module membership”) and then computes an “eigengene” for each sample and each module (corresponding to the first principal component of all genes in that module). We identified a total of 156 modules across six WGCNA runs (regions: cortex, amygdala; subregion: dACC, dlPFC, MeA, BLA; **Table S10**). We first tested for enrichment of module membership among differentially expressed genes (at p <0.005) identified for PTSD versus neurotypicals and MDD versus neurotypicals. There were 35 total modules enriched for genes implicated in either disorder (at FDR < 0.05, odds ratio, OR>1; PTSD: 22 modules, MDD: 22 modules, with 9 in common). We further tested for module eigengene associations with PTSD, MDD, and PTSD-specific diagnoses for convergent evidence implicating each module with each disorder, and annotated each module with the most significant gene ontology-enriched category (**Table S11**). In the cortex and its subregions, the strongest disorder-related module (Cortex.ME7) related to regulation of cell activation, a broad category encompassing many immune processes, associated with both PTSD (p=1.6e-25, **Table 3**) and MDD (p=3.3e-126) DEGs, with its eigengene further associated with these diagnoses at the subject-level (PTSD p=2.9e-4, MDD p=8.4e-6). The strongest disorder-related module in the amygdala (Amygdala.ME2) was specifically enriched with PTSD DEGs (p=2.97e-23) with its eigengene further associated with PTSD compared to controls (p=0.005). These analyses further highlight biological processes associated with PTSD and MDD using convergent approaches to traditional gene set enrichments of DEGs.

**Table 3:** Module-level associations to PTSD and MDD. Fisher’s exact test enrichment for DEGs (at p < 0.005) in PTSD (PTSD_Pval) and MDD (MDD_Pval) among module gene membership. Eigengene subject-level associations to PTSD vs control (PTSD_p), MDD vs control (MDD_p) and PTSD versus combined MDD + Control (onlyPTSD_p). The top gene ontology biological process is shown for module gene membership (GOBP_Description) with corresponding p-value (GOBP_pvalue).

| Module_Name | numGenes | DEG Enrichment |  | Eigengene Assoc. |  | Gene Ontology (BP) |  |
| --- | --- | --- | --- | --- | --- | --- | --- |
|  |  | PTSD | MDD | PTSD | MDD | Description | P-value |
| Cortex_ME2 | 1360 | 4.40E-03 | 4.66E-11 | 1.69E-01 | 2.58E-05 | synapse organization | 3.49E-14 |
| Cortex_ME7 | 620 | 1.57E-25 | 3.29E-126 | 2.85E-04 | 8.45E-06 | regulation of cell activation | 1.68E-42 |
| Cortex_ME18 | 125 | 3.61E-01 | 2.22E-03 | 3.03E-01 | 5.81E-04 | modulation of chemical synaptic transmission | 6.21E-07 |
| Cortex_ME30 | 56 | 7.94E-03 | 6.65E-01 | 3.86E-01 | 2.14E-01 | regulation of neurotransmitter receptor activity | 8.82E-05 |
| Cortex_ME31 | 55 | 1.70E-10 | 1.88E-01 | 3.22E-02 | 2.42E-01 | learning or memory | 1.37E-04 |
| Cortex_ME34 | 35 | 7.27E-03 | 1.00E+00 | 8.39E-02 | 3.82E-01 | response to cAMP | 1.55E-08 |
| dIPFC_ME3 | 1104 | 3.53E-01 | 1.12E-03 | 1.53E-01 | 2.78E-03 | regulation of synaptic plasticity | 6.77E-12 |
| dIPFC_ME6 | 502 | 7.77E-05 | 8.26E-02 | 2.35E-03 | 6.92E-03 | regulation of leukocyte activation | 5.16E-41 |
| dIPFC_ME11 | 299 | 6.12E-01 | 1.89E-06 | 1.93E-02 | 6.93E-04 | meiotic chromosome separation | 1.87E-04 |
| dIPFC_ME20 | 117 | 2.71E-15 | 4.44E-02 | 9.14E-03 | 3.23E-01 | UDP-N-acetylglucosamine metabolic process | 1.36E-04 |
| dIPFC_ME22 | 98 | 2.37E-03 | 6.32E-01 | 1.38E-01 | 8.25E-01 | response to estradiol | 4.15E-04 |
| dIPFC_ME24 | 84 | 1.61E-04 | 2.69E-03 | 5.41E-02 | 7.92E-02 | membrane depolarization during action potential | 7.82E-07 |
| dACC_ME3 | 1393 | 1.55E-03 | 1.61E-23 | 1.01E-02 | 1.84E-08 | modulation of chemical synaptic transmission | 2.95E-22 |
| dACC_ME5 | 747 | 3.60E-03 | 2.70E-06 | 2.20E-02 | 1.10E-03 | forebrain development | 6.78E-10 |
| dACC_ME7 | 702 | 2.66E-03 | 3.28E-84 | 1.26E-03 | 2.03E-05 | lymphocyte activation | 3.57E-42 |
| dACC_ME9 | 599 | 1.42E-08 | 6.02E-01 | 4.79E-01 | 4.68E-02 | modulation of chemical synaptic transmission | 2.01E-09 |
| dACC_ME12 | 197 | 1.03E-01 | 6.02E-04 | 8.96E-01 | 1.11E-01 | heart development | 1.48E-06 |
| dACC_ME13 | 177 | 6.56E-03 | 1.00E+00 | 6.39E-01 | 8.99E-02 | negative regulation of translation | 9.03E-05 |
| Amygdala_ME1 | 1035 | 1.90E-05 | 3.66E-04 | 2.49E-01 | 3.99E-02 | myelination | 2.73E-13 |
| Amygdala_ME2 | 904 | 2.97E-23 | 2.59E-03 | 4.70E-03 | 1.15E-01 | lymphocyte activation | 1.70E-38 |
| Amygdala_ME4 | 696 | 1.90E-01 | 3.67E-05 | 2.56E-01 | 1.12E-01 | modulation of chemical synaptic transmission | 5.27E-26 |
| Amygdala_ME19 | 47 | 9.15E-02 | 7.72E-03 | 7.88E-03 | 5.43E-03 | extracellular matrix constituent secretion | 3.05E-04 |
| Amygdala_ME21 | 39 | 8.55E-03 | 2.86E-01 | 1.49E-02 | 1.15E-02 | regulation of system process | 3.77E-03 |
| Amygdala_ME24 | 32 | 2.94E-01 | 2.60E-03 | 3.17E-01 | 1.09E-01 | formation of quadruple SL/U4/U5/U6 snRNP | 4.84E-07 |
| MeA_ME3 | 431 | 9.88E-12 | 5.72E-01 | 4.09E-02 | 5.13E-01 | modulation of chemical synaptic transmission | 7.17E-18 |
| MeA_ME4 | 375 | 3.63E-01 | 3.22E-04 | 2.60E-01 | 4.32E-02 | modulation of chemical synaptic transmission | 1.71E-28 |
| MeA_ME5 | 281 | 1.11E-01 | 3.82E-03 | 6.05E-02 | 5.99E-03 | regulation of ion transmembrane transport | 1.21E-07 |
| MeA_ME7 | 153 | 5.00E-03 | 2.78E-06 | 1.53E-01 | 7.61E-02 | extracellular structure organization | 2.98E-10 |
| MeA_ME9 | 86 | 6.34E-01 | 7.33E-04 | 4.35E-01 | 2.20E-03 | ameboid-type cell migration | 1.45E-04 |
| MeA_ME10 | 85 | 3.21E-01 | 1.65E-05 | 1.01E-01 | 4.65E-04 | regulation of membrane potential | 7.35E-06 |
| MeA_ME13 | 63 | 7.22E-21 | 2.34E-26 | 5.51E-03 | 7.62E-04 | extracellular matrix organization | 1.36E-18 |
| BLA_ME2 | 1300 | 3.25E-09 | 5.09E-01 | 1.32E-02 | 9.98E-02 | lymphocyte activation | 1.81E-30 |
| BLA_ME12 | 199 | 4.08E-03 | 6.60E-02 | 2.52E-02 | 1.37E-01 | homophilic cell adhesion via plasma membrane adhesion molecules | 4.41E-06 |
| BLA_ME13 | 176 | 2.36E-07 | 6.47E-01 | 3.85E-03 | 5.21E-01 | locomotory behavior | 1.98E-06 |
| BLA_ME20 | 37 | 5.31E-02 | 1.13E-08 | 6.37E-02 | 1.17E-02 | spliceosomal tri-snRNP complex assembly | 9.98E-08 |

## Discussion

In this study we performed RNA-seq on homogenate tissue from two frontal cortical subregions (dlPFC and dACC) and two amygdala subregions (MeA and BLA) using the largest available sample of postmortem human brain tissue from individuals with PTSD available. Importantly, this cohort contained individuals with both primary and secondary diagnoses of PTSD (62.6% of these comorbid with MDD), individuals with MDD, but no PTSD diagnosis, and neurotypical control donors with no psychiatric diagnoses. We compared gene expression signatures across these groups to identify shared and divergent molecular signatures associated with PTSD and MDD in cortico-amygdala circuits. Differential expression analysis identified a small number of genes associated specifically with PTSD diagnosis, which were predominantly identified in the cortical regions.

We observed consistent down-regulation of *CORT*, a marker gene for cortistatin-positive interneurons across all four subregions in individuals with PTSD, and cell-specific expression analysis (CSEA) of differentially expressed genes confirmed enrichment of these DEGs in cortistatin-positive interneurons in PTSD. We confirmed co-expression of *CORT* with a broad interneuron marker (*GAD2*) as well as additional interneuron markers that were differentially expressed in PTSD (*SST* and *CRHBP*) using single molecule fluorescence in situ hybridization (smFISH) in amygdala of the human brain. Weighted gene coexpression network analyses (WGCNA) further implicated interneuron and immune system function in line with the gene set enrichment results applied directly to PTSD DEGs. smFISH in sections of the postmortem human amygdala confirmed that expression of *CORT, SST, and CRHBP* transcripts are localized to GABAergic inhibitory neurons. Downregulation of these transcripts in our study lends further support to the hypothesis that interneuron dysfunction is mechanistically associated with PTSD (23). There is strong evidence that GABAergic interneurons control neural activity and synaptic plasticity in cortico-amygdala circuits to regulate fear-related behaviors in preclinical animal models that are relevant for PTSD and other trauma-related disorders (24). For example, within the basolateral amygdala (BLA), the activity of excitatory cells that project to the frontal cortex is under tight regulation by local GABAergic inhibitory neurons (26). Inhibitory control over these excitatory projections tightly controls top down negative feedback regulation of the BLA in the expression and extinction of fear, which is highly relevant for PTSD and other trauma-related disorders (27; 28).

Decreased expression of genes included immune-related gene ontology sets were associated with PTSD diagnosis in both cortical and amygdala brain regions (**Figure 3A**). Supporting the notion of immune signaling involvement in PTSD, CSEA in mouse and enrichment analyses using human single nucleus RNA sequencing (snRNA-seq) data demonstrated enrichment of these DEGs in mouse and human microglia profiles (63; 64). Genes with decreased expression in MDD donors compared to neurotypical controls showed analogous enrichment of immune-related processes using both gene set enrichment analysis and CSEA; however, there were no enrichments for the immune-related gene sets when contrasting PTSD and MDD, suggesting decreased expression of immune processes and microglia involvement are not specific to PTSD. The direction of dysregulation - decreased expression of immune and/or microglia-related genes - may be considered surprising considering that higher levels of pre-trauma C-reactive protein, a marker of blood inflammation, predicted elevated PTSD symptoms after trauma, and elevated levels of selected markers of low-grade blood inflammation have been reported in a meta-analysis of PTSD studies (76; 77). However, over time, and with repeated exposure to trauma or as the result of chronic stress as a sequelae of the trauma, immune function can be dysregulated in a myriad of ways, with both neuronal and peripheral systems attempting to compensate for immune activation and increased inflammation (35; 78–80). Moreover, we identified that a number of the genes included in the gene ontology sets related to immune signaling encode proteins with known immunosuppressive activity, which could also explain the somewhat paradoxical finding of decreased expression of immune-related genes. For example, in the immune-related regulation of cell activity category, we identified 13 member PTSD DEGs - and of these 13, 7 (bolded) are suggested to have potential immunosuppressive activity in the human (*IL1RL2/**DPP4/**IGFBP2/**TGFBR2**/TAC1/MDK/**CD4**/PTPN6/TESPA1/**IGF1**/**ITGAM**/**TYROBP**/**ITG B2***) (81).

We further ran a number of analyses to better untangle the gene expression effects that were selectively associated with PTSD, and not MDD. In general, we identified more DEGs for MDD than PTSD, particularly in the cortex (which were primarily driven by the dACC). However, gene expression differences were highly concordant between the two diagnoses, with all significant DEGs showing the same directionality of effects (ie log2 fold changes) in both diagnoses. We nevertheless identified a small number of marginally significant DEGs when directly comparing patients with PTSD to MDD in each brain subregion. Those DEGs more highly expressed in PTSD compared to MDD in cortex were enriched for glutamatergic synapses (driven by the dACC) and those DEGs more highly expressed in MDD compared to PTSD were enriched for neuronal activity in amygdala (driven by the BLA). These differences between the two diagnoses were further magnified in WGCNA analyses, where seven (potentially overlapping) modules showed very PTSD-specific enrichment (Cortex_ME31, dlPFC_ME20, dACC_ME9, Amygdala_ME2, MeA_ME3, BLA_ME2, BLA_ME13) and five modules showed very MDD-specific enrichment (dlPFC_ME11, dACC_ME3, dACC_ME7, MeA_ME9, BLA_ME20).

Differentially expressed genes (DEGs) from a recent RNA-seq study of human postmortem PTSD tissue(70), which used a partially overlapping set of donors (see below) provides support for top DEGs identified here. For example, within our combined cortical PTSD analyses (see Figure 1), 6 of the 7 most robustly affected transcripts comparing PTSD versus controls (*CORT, HDAC4, CRHBP, ADAMTS2, FBXO9, APOC1*) were directionally consistent and at least marginally significant in this previous dataset. These genes further showed decreased expression in MDD versus controls in the present study, with at least marginal significance, suggesting that these particular findings may be related to shared pathophysiological changes accompanying PTSD and MDD versus neurotypical gene expression. A key up-regulated gene identified in Girgenti et al., *ELK1*, was significantly up-regulated in both cortical regions in the present study, and the somatostatin gene (*SST*), identified as robustly down-regulated in several regions of the cortex in Girgenti et al., was in the top ten of all down-regulated transcripts in both dlPFC and dACC here. *ADAMTS2*, the second highest upregulated DEG in the combined cortical sample, was the top up-regulated gene in the dACC and the third most up-regulated in the dlPFC in Girgenti et al., 2020 (82). Both studies further found enrichment of PTSD-associated DEGs related to interneurons and their molecular functions.

In addition to these common elements, the present results extend previous findings Girgenti et al., 2020 in several key areas. First, this study extended the search for differential gene expression beyond the cortex and into the amygdala, a relatively under-studied brain area in postmortem human brain research with high relevance to PTSD. Second, we provide compelling evidence implicating *decreased* expression of immune-related genes and associated processes in PTSD and MDD compared to neurotypical controls. This is an important observation because it runs counter to the most likely expectations for directionality. We further refined interneuron enrichment more specifically to *CORT*-positive interneurons, which we subsequently validated with RNAscope. The cell type analyses in the present study provide direct evidence of these enrichments by interrogating DEGs directly against cell type-specific genes from both human and mouse studies, complementing the the indirect strategy taken by Girgenti et al of first identifying genes in discrete co-expressed modules, and then associating those genes with both PTSD DEGs and cell type-specific genes separately (such that different genes capturing the cell type versus PTSD signal in the same module). Third, we believe our >2-fold increased sample size in all diagnostic groups (total N=325 versus N=143) - obtained from a single postmortem brain collection under identical sample ascertainment and inclusion criteria - refined several of the clinical associations identified by Girgenti et al. We identified more similarities than differences between PTSD and MDD and replicated this finding across both amygdala and cortical subregions, with far less sex-specific diagnosis-associated signal. Contributing to both the similarities and differences between the two studies was the fact that 77 donors were shared across both studies, although the two studies used different hemispheres, independent dissections and RNA extractions, as well as different data analysis pipelines. Over half (53.8%) of the donors in Girgenti et al., and a quarter (23.7%) of the donors overlapped with the present study.

Therefore, it might seem counter-intuitive that we identified many fewer DEGs in this much larger study, particularly with overlapping donors. We believe these differences can be accounted for by our more conservative statistical analyses - including modeling both diagnostic groups in a single statistical model against the neurotypical group that further accounted for robust observed and latent confounders. The differential expression models in Girgenti et al only adjusted for age, RIN, PMI, and race, and lack of accounting for sequencing-derived RNA quality metrics and other latent confounders, which can greatly increase false positive rates in human postmortem brain gene expression studies (48). For example, this less comprehensive statistical model applied to our larger dataset resulted in 1,243 DEGs in DLPFC, 1,719 DEGs in dACC, 1,813 DEGs in BLA and 10,283 DEGs in MeA for PTSD at FDR < 0.05, many more genes than reported here. There has been some debate regarding the optimal methods of latent variable correction in these postmortem studies (and the potential for “over-correction”) (83). A major analytic element of the present investigation was the use of quality surrogate variable (qSV) analysis to identify and correct for expressed sequences that are particularly prone to degrade in human post-mortem brain (48). The qSVs utilized here were defined from the top 1000 degradation-susceptible expressed regions generated from independent time course experiments. Dropping the qSVs from our main analyses resulted in 209 DEGs in DLPFC, 43 DEGs in dACC, 62 DEGs in BLA and 1,054 DEGs in MeA (at FDR < 0.05) for PTSD in the present dataset. Potentially, if there are sex or disease-associated interactions related to gene transcript degradation, use of qSV may have limited the emergence of these genes as DEGs and contributed to the differences between the present findings and those discussed in Girgenti et al. Similarly, in MDD, the most prominent previous report of differential gene expression actually identified no DEGs when correcting for multiple testing via the FDR (from the supplementary tables included in that manuscript) (84) making it difficult to assess replication of our DEGs using previously-published datasets. While these issues may seem rather nuanced, they nevertheless have important consequences on identifying DEGs in human postmortem RNA-seq datasets, and require careful consideration in past and future work.

Overall, these analyses of the largest postmortem brain cohort of patients with PTSD and MDD to date highlight the sub-population of cortistatin-expressing interneurons as having potential functional significance in PTSD, and provide evidence for dysregulated neuroinflammation and neuroimmune signaling in MDD and PTSD pathophysiology.

## Supporting information

Differential Expression Statistics

## Data availability

All code and figures associated with this manuscript are available through GitHub: https://github.com/LieberInstitute/LIBD_VA_PTSD_RNAseq_4Region. All raw and processed data will be made available through a Globus endpoint.

## Acknowledgements

The authors would like to express their gratitude to our colleagues whose tireless efforts have led to the donation of postmortem tissue to advance these studies: the Office of the Chief Medical Examiner of the District of Columbia, the Office of the Chief Medical Examiner for Northern Virginia, Fairfax Virginia, the Office of the Chief Medical Examiner of the State of Maryland, Baltimore Maryland, the Office of the Chief Medical Examiner for Kalamazoo County, Kalamazoo Michigan, the University of North Dakota School of Medicine Department of Pathology, Forensic Pathology Center, Grand Forks North Dakota, and the Santa Clara County Office of the Chief Medical Examiner, San Jose California. This work was supported with resources and use of facilities at the VA Connecticut Health Care System, West Haven, CT, Central Texas Veterans Health Care System, Temple, TX, Durham VA Healthcare System, Durham NC, VA San Diego Healthcare System, La Jolla, CA, VA Boston Healthcare System, Boston, MA, USA and the National Center for PTSD, U.S. Department of Veterans Affairs. The views expressed here are those of the authors and do not necessarily reflect the position or policy of the Department of Veterans Affairs (VA) or the U.S. government. We also would like to acknowledge the contributions of Amy Deep-Soboslay and Llewellyn B. Bigelow, MD for their diagnostic expertise. Finally, we are indebted to the generosity of the families of the decedents, who donated the brain tissue used in these studies. We also would like to acknowledge the contributions of Llewellyn B. Bigelow, MD for his diagnostic expertise, and Daniel R. Weinberger for providing constructive commentary and editing of the manuscript. Finally, we are indebted to the generosity of the families of the decedents, who donated the brain tissue used in these studies.

## Supplementary Figures

**Figure S1:**
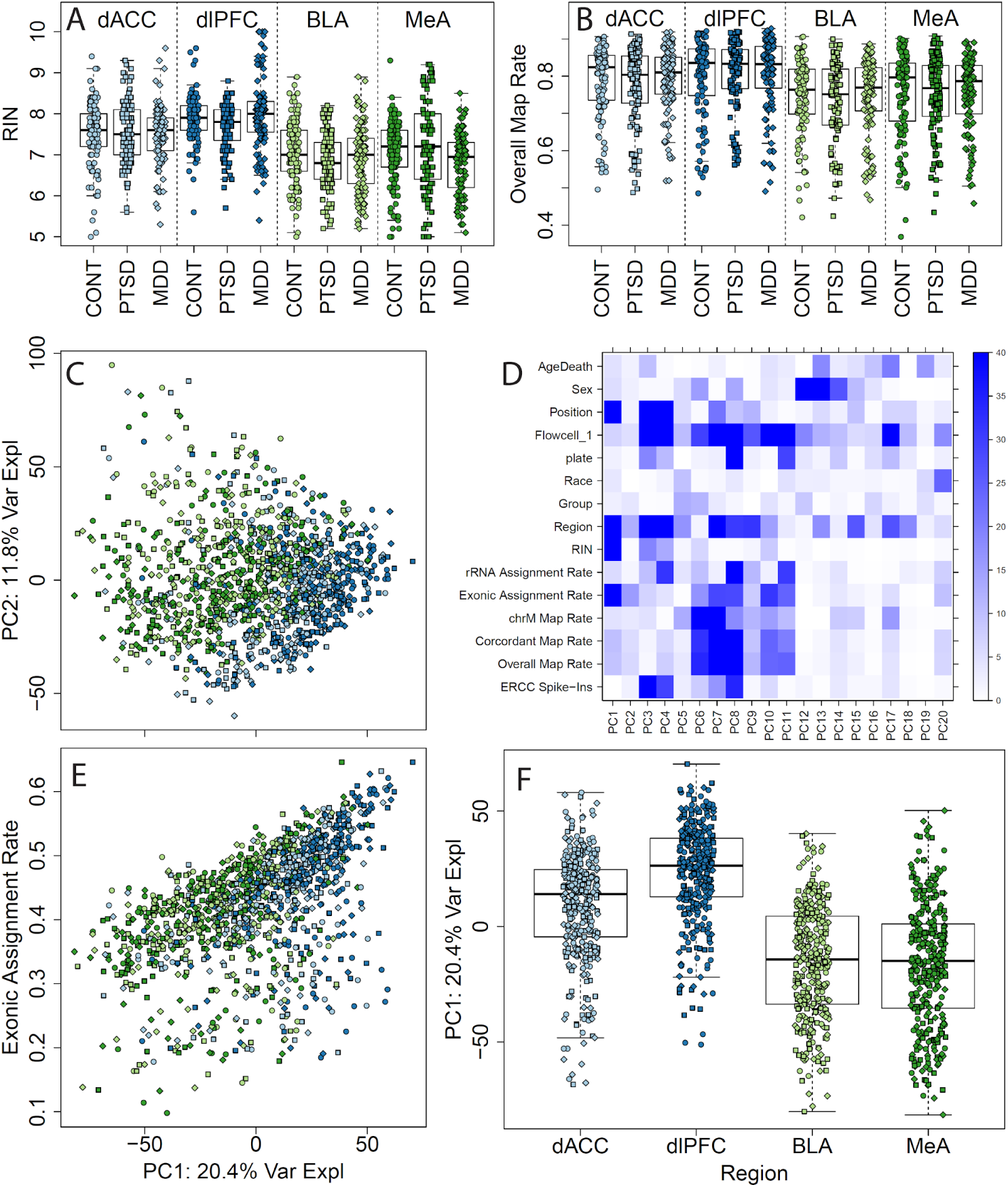
RNA quality and sequencing metrics. (A) RNA integrity numbers (RINs) and (B) overall RNA-seq read mapping rates across brain regions and diagnosis groups. (C) Principal component (PC) 1 versus 2 shows differences by brain region. (D) Associating observed clinical and technical variables with gene expression PCs, colors are negative log10 p-values from linear regression (either single terms for continuous or binary variables and ANOVA group p-values for categorical variables). (E) Exonic assignment rate (i.e. the fraction of aligned reads that were assigned to genes during counting) and (F) brain region, particularly cortex versus amygdala, associates with PC1 as well.

**Figure S2:**
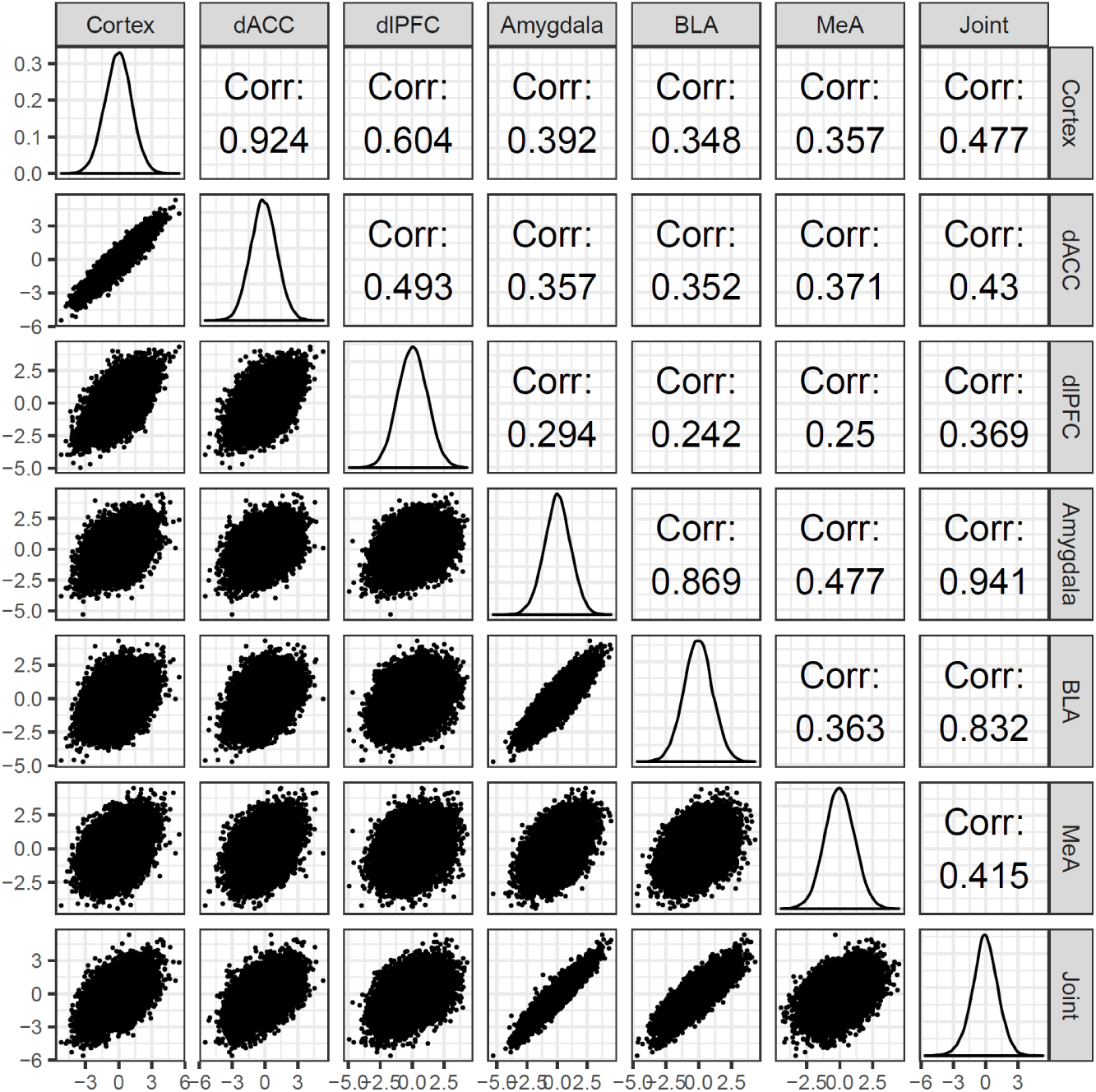
High gene-level correlations between PTSD effects across different subsets of samples, both within and across brain subregions.

**Figure S3:**
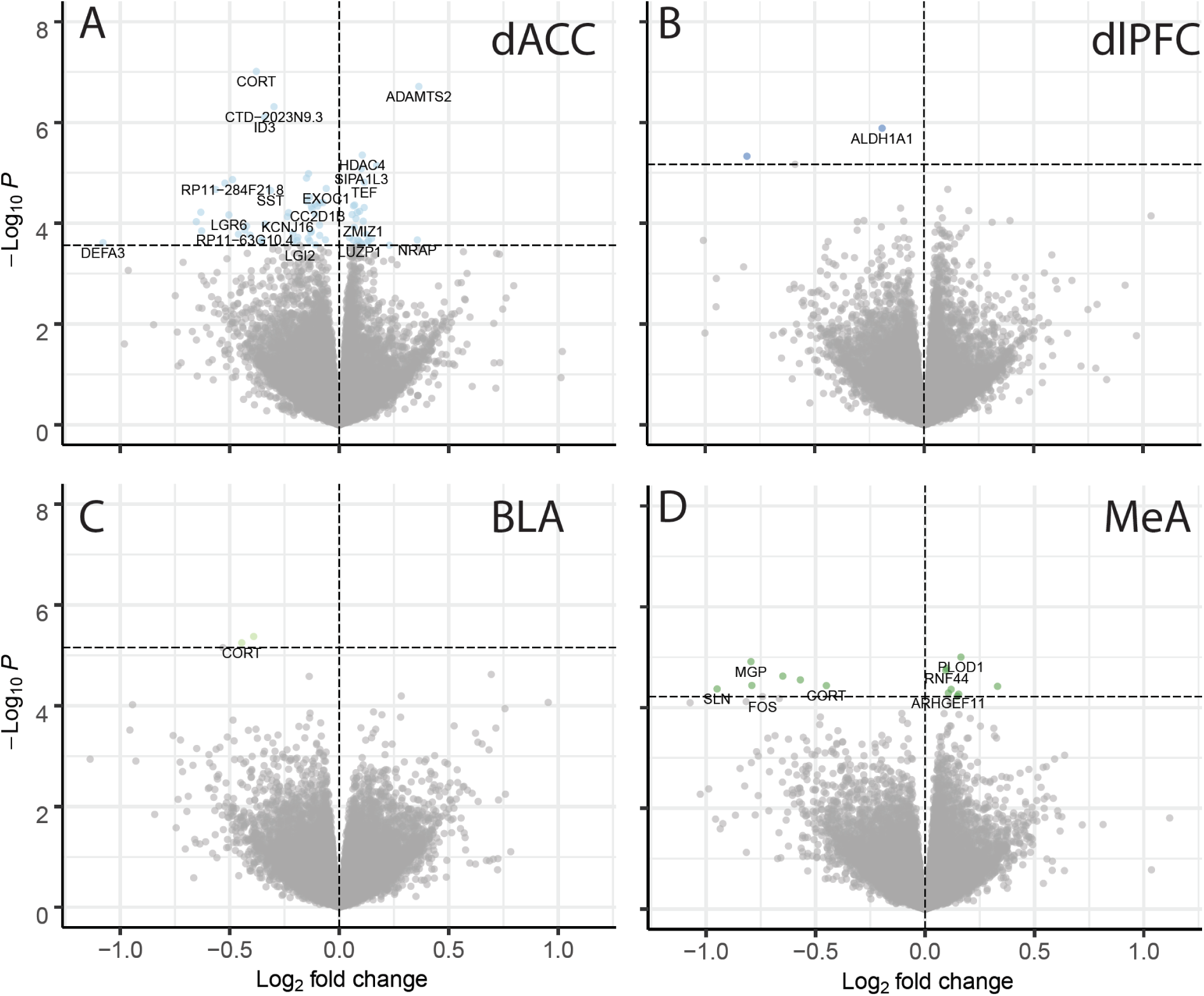
Volcano plots by brain subregions, for (A) dACC and (B) dlPFC in the cortex and the (C) BLA and (D) MeA in the amygdala. Horizontal lines represent p-values that control FDR < 0.1.

**Figure S4:**
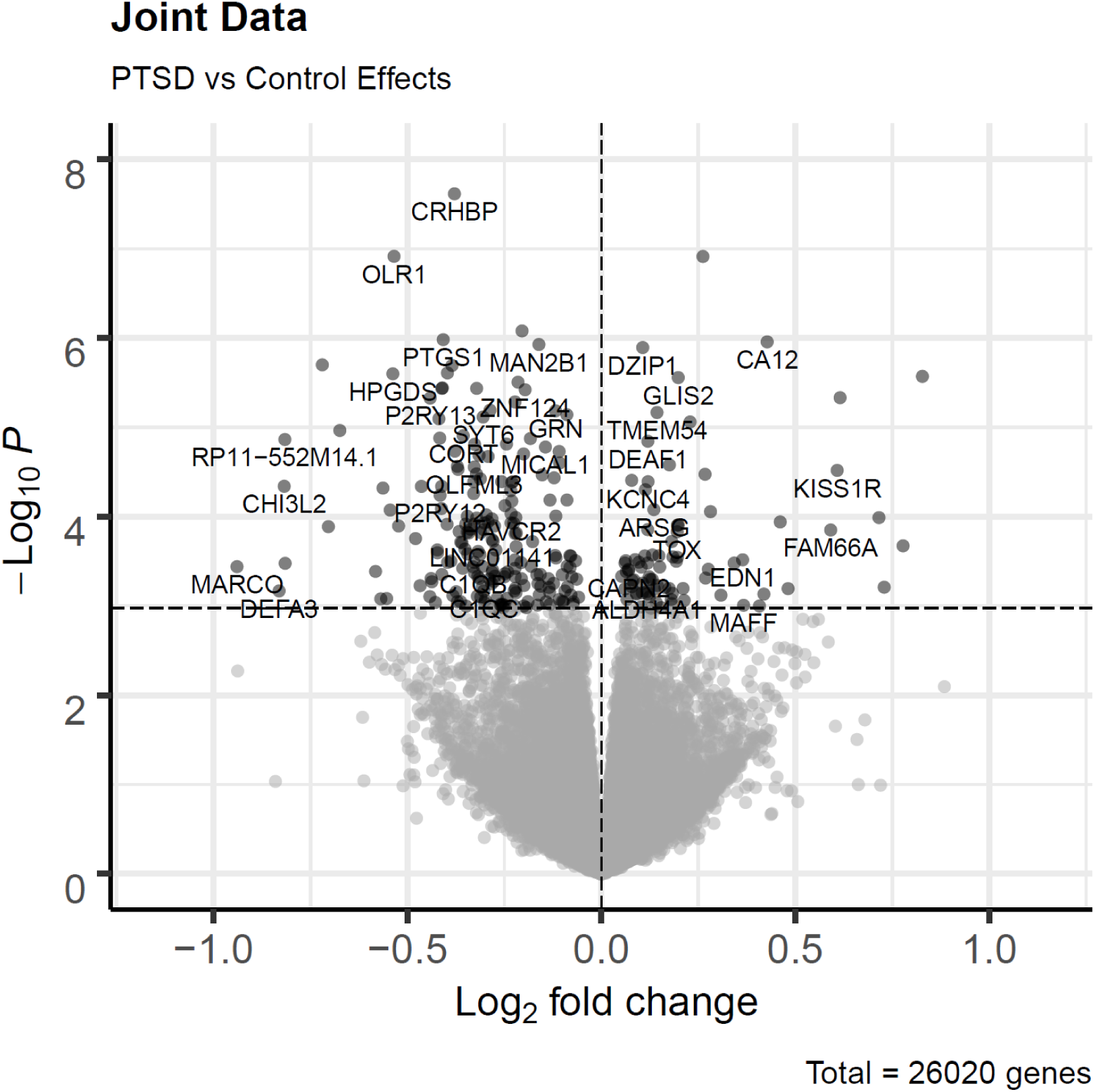
volcano plot across all regions.

**Figure S5:**
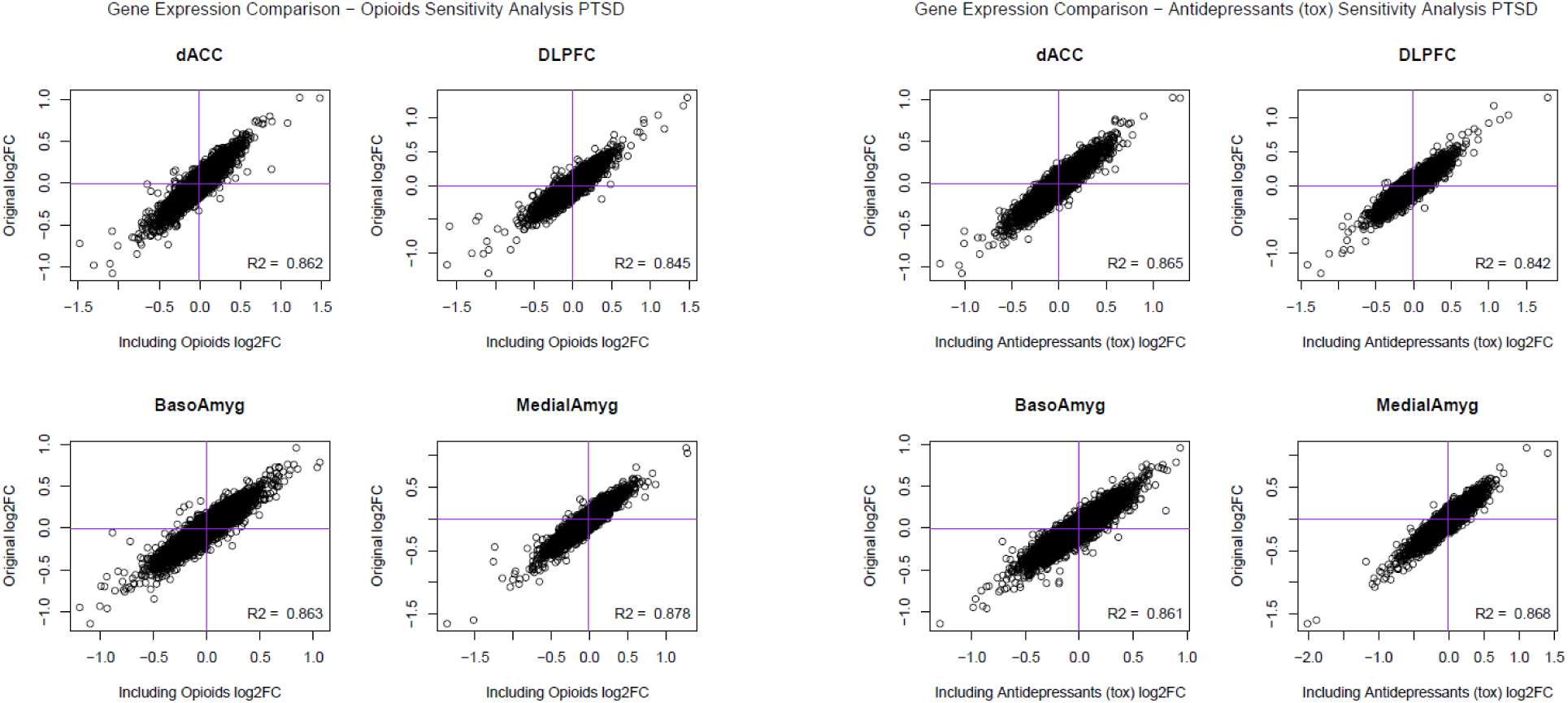
Sensitivity analyses for PTSD effects further adjusting for A) opioid and B) antidepressant exposures, in addition to all other considered observed and latent confounders described in the Methods section.

**Figure S6:**
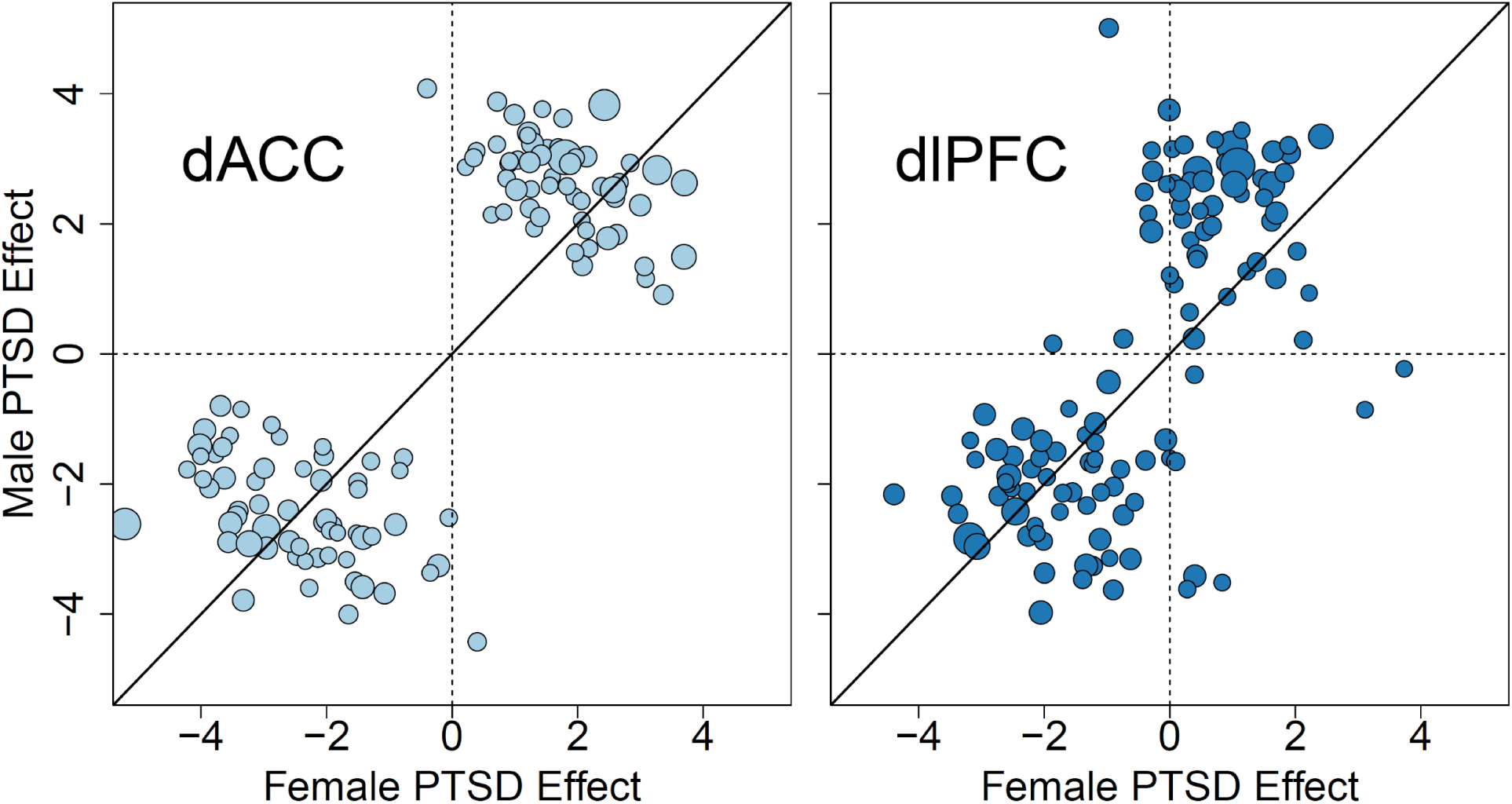
Differences in PTSD effects across the two sexes in the two cortical brain regions. Effects shown are T-statistics.

**Figure S7:**
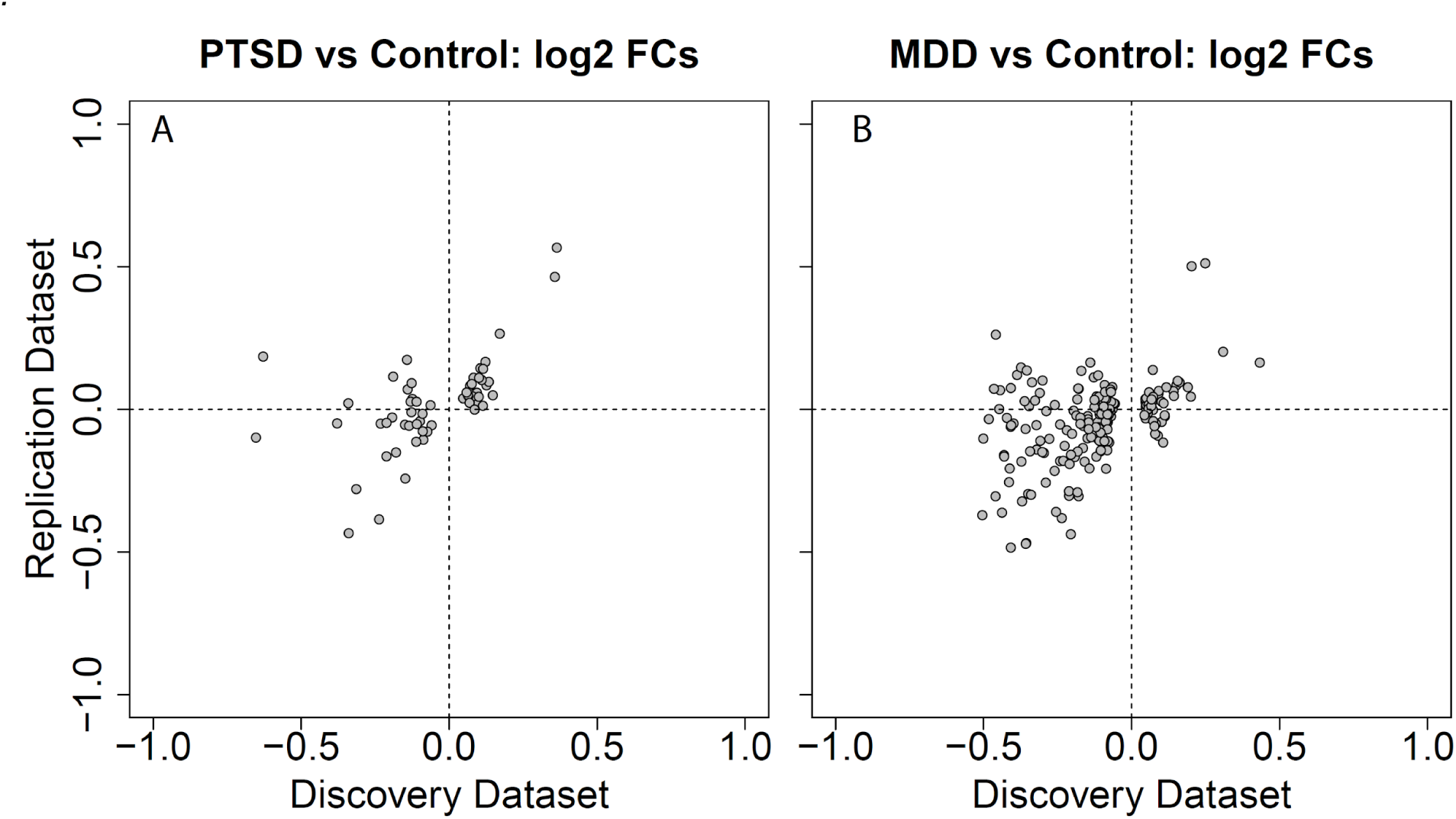
Replication of dACC effects with Girgenti et al 2020 (70). (A) PTSD log2 fold changes and (B) MDD log2 fold changes for DEGs identified here at FDR < 0.1.

**Figure S8:**
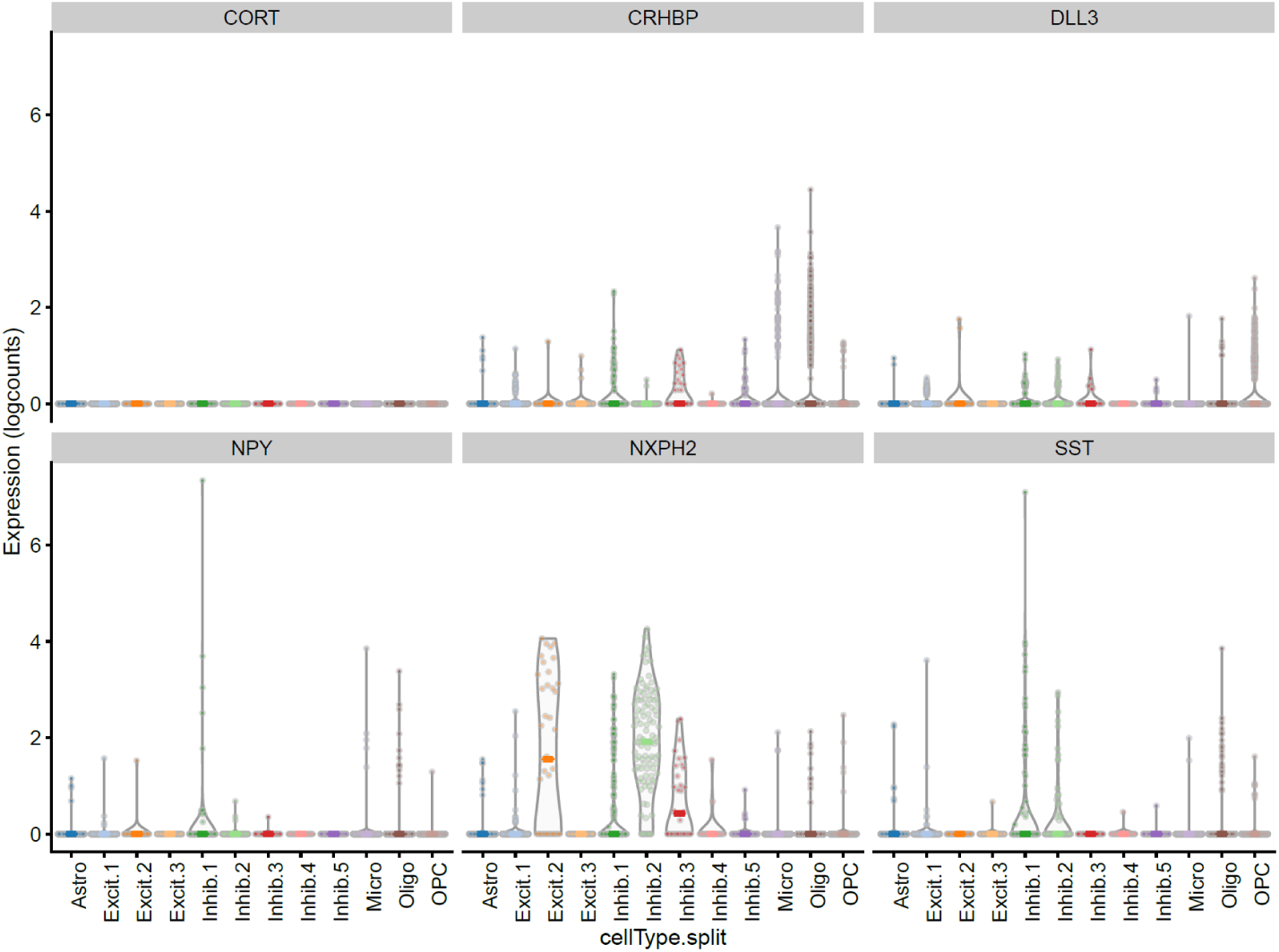
expression of interneuron DEGs in amygdala snRNA-seq data, including CORT

**Figure S9:**
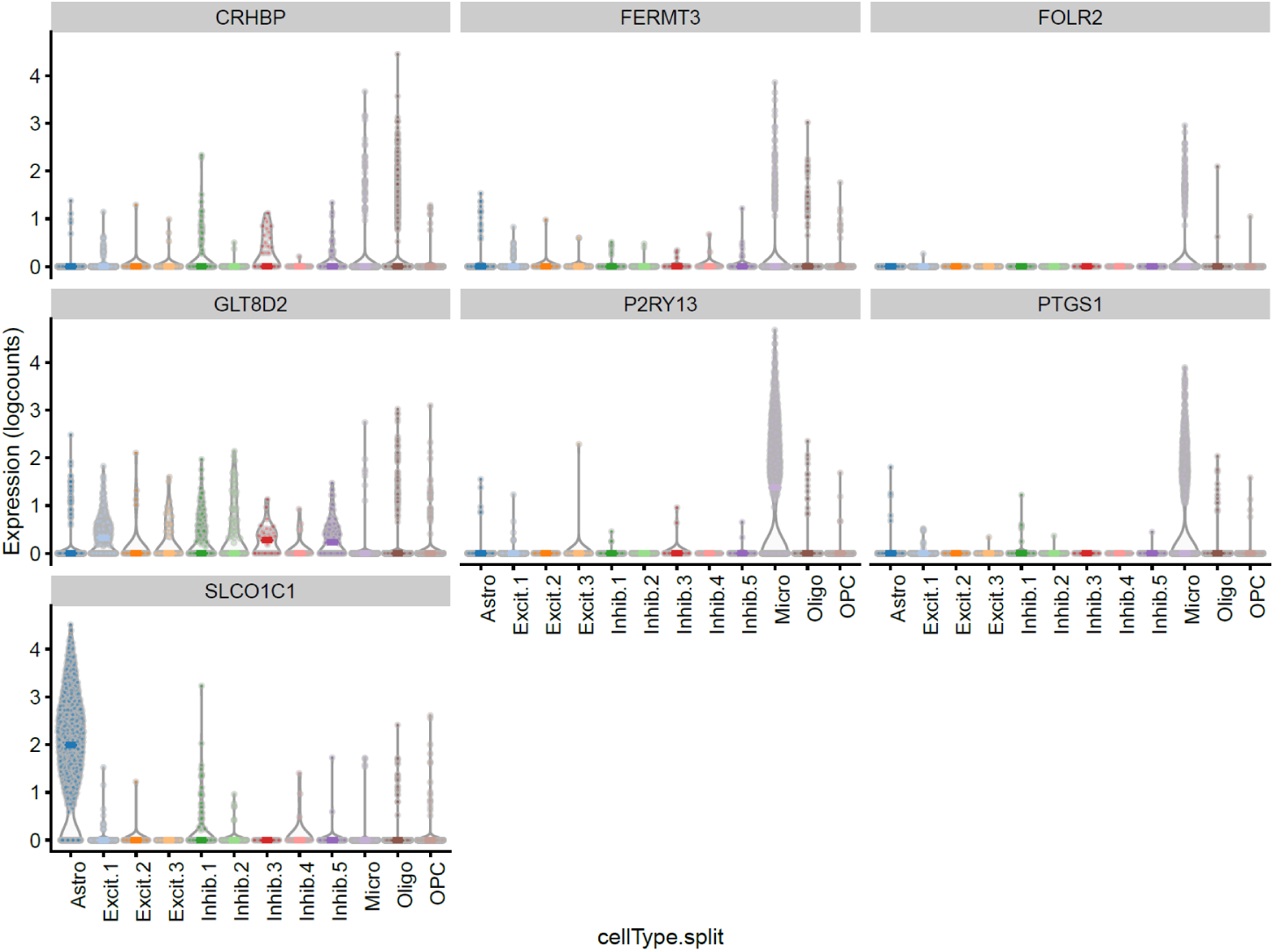
expression of immune-related DEGs in amygdala snRNA-seq data

**Figure S10:**
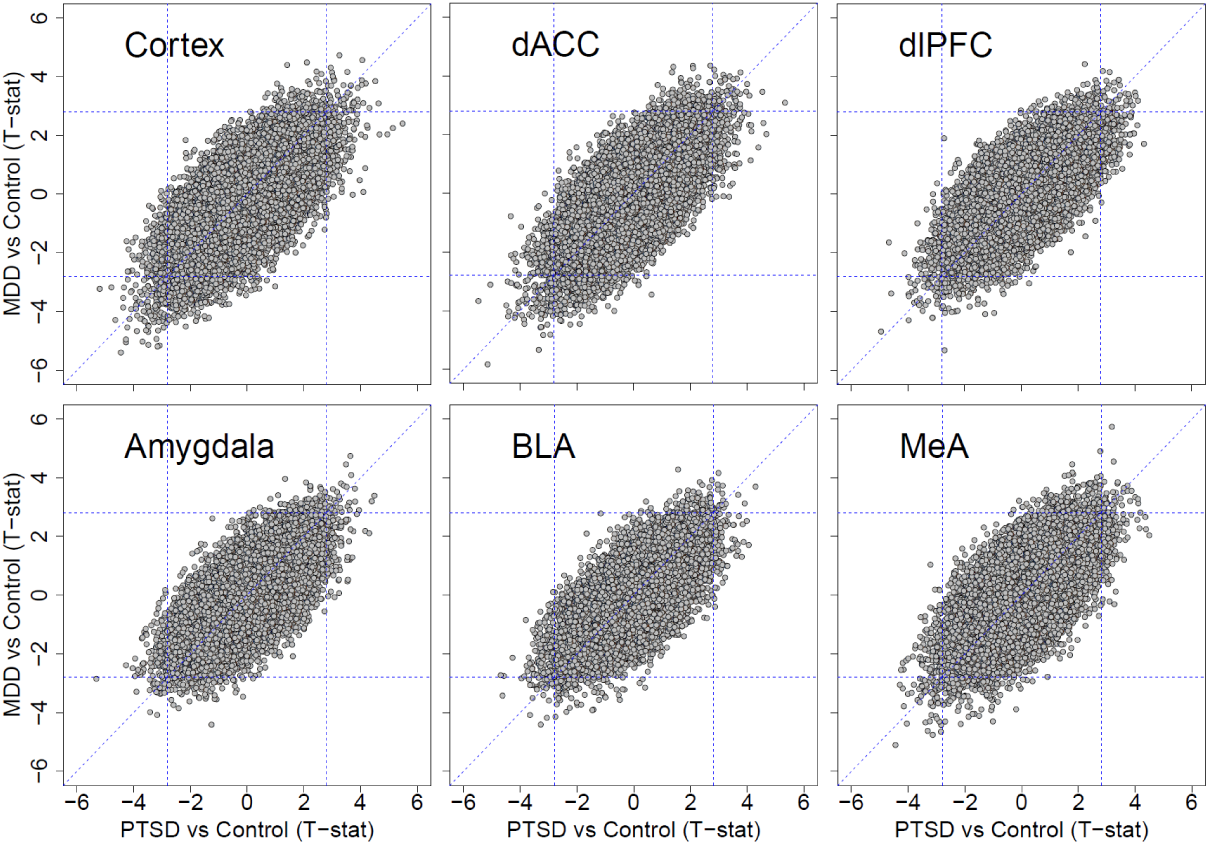
High correlation between PTSD and MDD effects across brain regions

**Figure S11:**
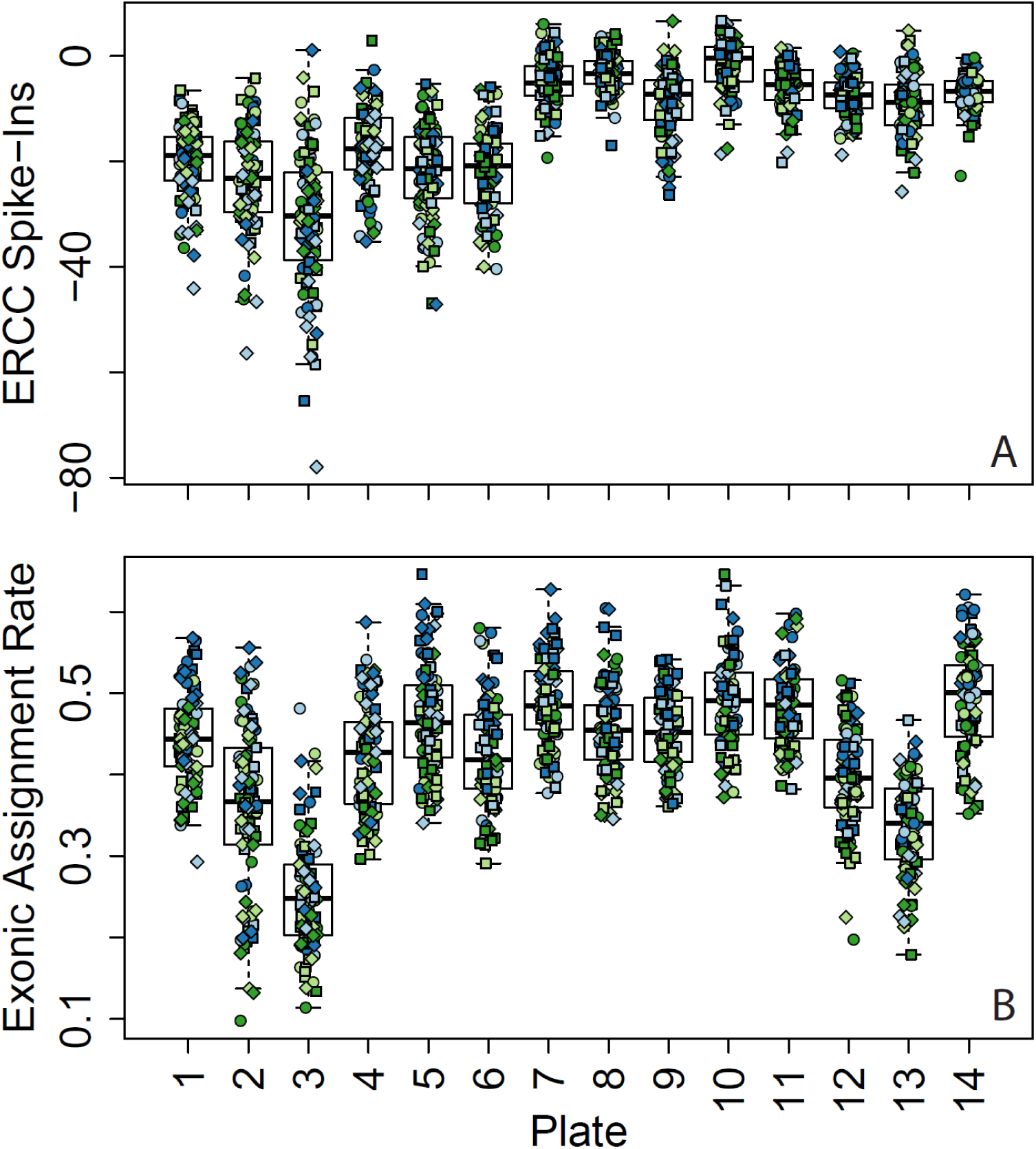
Sequencing metrics by processing plate. (A) ERCC Spike-in bias and (B) exonic mapping rates across the 14 processing plates. Colors indicate brain region and shapes indicate diagnostic groups (using same coloring as Figure 1)

## Notes

### Competing Interest Statement

The authors have declared no competing interest.

### Summary of Updates

One author wished to be removed from the manuscript

